# Intracellular *Staphylococcus aureus* perturbs the host cell Ca^2+^-homeostasis to promote cell death

**DOI:** 10.1101/2020.08.20.260471

**Authors:** Kathrin Stelzner, Ann-Cathrin Winkler, Liang Chunguang, Carsten P. Ade, Thomas Dandekar, Martin J. Fraunholz, Thomas Rudel

**Author notes:** Address correspondence to Thomas Rudel. Ann-Cathrin Winkler and Liang Chunguang contributed equally to this work.

## Abstract

The opportunistic human pathogen *Staphylococcus aureus* causes serious infectious diseases ranging from superficial skin and soft tissue infections to necrotizing pneumonia and sepsis. While classically regarded as extracellular pathogen, *S. aureus* is able to invade and survive within human cells. Host cell exit is associated with cell death, tissue destruction and spread of infection. The exact molecular mechanism employed by *S. aureus* to escape the host cell is still unclear. In this study, we performed a genome-wide shRNA screen and identified the calcium signaling pathway to be involved in intracellular infection. *S. aureus* induced a massive cytosolic Ca^2+^-increase in epithelial host cells after invasion and intracellular replication of the pathogen. This was paralleled by decrease in endoplasmic reticulum Ca^2+^-concentration. Additionally, calcium ions from the extracellular space contributed to the cytosolic Ca^2+^-increase. As a consequence, we observed that the cytoplasmic Ca^2+^-rise led to increase in mitochondrial Ca^2+^-concentration, the activation of calpains and caspases and eventually to cell lysis of *S. aureus*-infected cells. Our study therefore suggests that intracellular *S. aureus* disturbs the host cell Ca^2+^-homeostasis and induces cytoplasmic Ca^2+^-overload, which results in both apoptotic and necrotic cell death in parallel or succession.

**Importance:** Despite being regarded as an extracellular bacterium, the pathogen *Staphylococcus aureus* can invade and survive within human cells. The intracellular niche is considered as hide-out from the host immune system and antibiotic treatment and allows bacterial proliferation. Subsequently, the intracellular bacterium induces host cell death, which may facilitate spread of infection and tissue destruction. So far, host cell factors exploited by intracellular *S. aureus* to promote cell death are only poorly characterized. We performed a genome-wide screen and found the calcium signaling pathway to play a role in *S. aureus* invasion and cytotoxicity. The intracellular bacterium induces a cytoplasmic and mitochondrial Ca^2+^-overload, which results in host cell death. Thus, this study firstly showed how an intracellular bacterium perturbs the host cell Ca^2+^-homeostasis.

## Introduction

*Staphylococcus aureus* is a Gram-positive pathogen frequently colonizing part of the human microflora (1). Despite its common mode of asymptomatic colonization in the healthy population, immunocompromised patients often suffer from serious infection outcomes, which vary from superficial skin lesions to deep-seated abscesses and life-threatening sepsis and are often caused by the endogenous carriage strain (2, 3). *S. aureus* infections thus lead to high morbidity and mortality and healthcare-associated costs were exacerbated by *S. aureus* isolates with resistance against many antibiotics (4).

*S. aureus* has been demonstrated to invade many mammalian cell types, such as epithelial and endothelial cells, osteoblasts, fibroblasts or keratinocytes. The pathogen has been shown to survive inside host cells for extended periods of time, not only *in vitro* (e.g. 5, 6–10), but also *in vivo* where intracellular *S. aureus* has been detected in tissue samples and phagocytic cells (11–15). Intracellularity of *S. aureus* is associated with chronic or relapsing staphylococcal infections as well as resistance to the host immune system and antibiotic treatment.

*S. aureus* adherence and subsequent uptake by non-professional phagocytes is facilitated by several bacterial adhesins, which interact with extracellular matrix proteins or specific host receptors (16, 17). Translocation from the bacteria-containing vacuole to the host cell cytoplasm is required for intracellular replication of the pathogen in, for instance, epithelial cells (18). Infection of cultured cells with *S. aureus* often elicits severe cytotoxicity that is believed to be the cause of pathologies originating from infection-induced tissue destruction (3, 19). Various rounds of *S. aureus* internalization and escape from host cells may contribute to tissue and membrane barrier destruction, spread of infection, immune evasion and inflammation (20–22).

Previous work has demonstrated a fundamental role of staphylococcal virulence factors in cellular cytotoxicity, directly damaging the cell and causing apoptotic and/or necrotic types of cell death (23–25). Besides, *S. aureus* invasion of host cells was shown to be required to activate cell death mechanisms (26–31). Very little is known about the cellular signaling involved in staphylococcal intracellular infection. To date, studies of host cell death on non-professional phagocytes infected with *S. aureus* provide conflicting results. Hallmarks of apoptosis, such as DNA fragmentation, cell contraction and/or activation of caspases, were reported in *S. aureus*-infected epithelial cells (26, 28, 32, 33). In bovine mammary epithelial (MAC-T) cells the extrinsic apoptosis pathway involving inflammatory cytokines and caspase 8 and 3 was activated by intracellular *S. aureus* (34). In contrast, induction of autophagy but not activation of caspases was shown to be essential for *S. aureus* cytotoxicity (35). Details on how intracellular *S. aureus* interferes with cell death signaling remain to be elucidated.

Ca^2+^-signaling has been implicated in various steps of bacterial infection and several bacterial toxins are known to induce Ca^2+^-fluxes in host cells by the formation of pores in the plasma membrane and/or activation of Ca^2+^-channels (36). Due to its low cytosolic concentration, calcium ions act as second messengers controlling diverse cellular functions, such as cell survival and cell death (37–40). Increases in cytosolic Ca^2+^-levels originate from uptake of extracellular Ca^2+^ by plasma membrane Ca^2+^- channels or by extrusion of calcium ions from internal stores. Store-operated calcium entry (SOCE) represents the predominant Ca^2+^-entry mechanism in non-excitable cells. Here, Ca^2+^-depletion from the ER activates Ca^2+^-entry across the plasma membrane and thus refills internal Ca^2+^-stores (41). For instance, regulated Ca^2+^ elevations are involved in cell signaling during the intrinsic apoptotic pathway and local Ca^2+^ messages at the ER-mitochondria interface participate in early apoptosis. A family of cysteine proteases called calpains also plays a role in Ca^2+^-induced apoptosis (42). The Ca^2+^-sensitive proteases cleave several members of the caspase and BCL2 family to regulate apoptotic cell death. Further, high concentration of mitochondrial Ca^2+^ activates mitochondrial permeability transition (MPT)-driven necrosis (43, 44). Disruption of ER Ca^2+^ homeostasis may result in ER stress and unfolded protein response (UPR) eventually leading to cell death (45). When stress results in cellular Ca^2+^-overload or loss of Ca^2+^ homeostatic control, Ca^2+^ functions also as executer of cell death (38).

In this study, we investigated how intracellular *S. aureus* interferes with host cell Ca^2+^-signaling. A Ca^2+^-reporter cell line and live cell imaging was used to monitor Ca^2+^-fluxes in single cells infected with the pathogen. Our results indicate that the intracellular bacterium perturbs the host cell Ca^2+^-homeostasis to induce cell death.

## Results

### An unbiased shRNA screen identifies calcium-associated genes and pathways in *S. aureus* invasion and cytotoxicity

We performed a whole genome RNA interference (RNAi) screen using small-hairpin RNAs (shRNAs) to identify host cell genes involved in *S. aureus* invasion and cytotoxicity (Figure 1A). HeLa cells were transduced with a lentiviral shRNA library, thereby generating a pool of host cells, in which genes were knocked-down by RNAi. The used shRNA library comprised 52,093 different shRNAs targeting 11,226 different host genes. This pool of HeLa cells (“input”) was infected with *S. aureus* 6850, a strongly cytotoxic and cell invasive strain (46), expressing red-fluorescent protein (mRFP). After an infection pulse of 1 h, extracellular bacteria were removed by treatment with the staphylolytic protease lysostaphin to ensure that cytotoxicity was not exerted by residual bacteria in the tissue culture medium. Uninfected and infected cell populations were FACS-sorted 4.5 hours post-infection (Figure S1A). DNA was prepared from the “input” pool as well as from the “uninfected” and “infected” cells and individual shRNA frequencies were determined by next-generation sequencing. Quality controls and data normalization were performed and a principal component analysis (PCA) demonstrated the validity of the datasets (Figure S1B and C). Selective enrichment of shRNAs retrieved from “uninfected” and “infected” cells were compared to the input cell pool. We thereby quantified the differential shRNAs (False discovery rate (FDR) adjusted *P*-value < 0.001; Log2FC 1.5), of which 79 were enriched and 217 were depleted in the “uninfected” sample (Table S1), indicative of shRNAs that negatively and positively regulate the infection process, respectively. In the “infected” sample 233 shRNAs were enriched and 311 were depleted (Table S1), reminiscent of shRNAs that negatively and positively regulate infection-induced cell death. Genes targeted by shRNAs with log2FC 3 (Invasion) or 3.5 (Cytotoxicity), respectively, are illustrated in heatmaps (Figure S1D and E).

**Figure 1.**
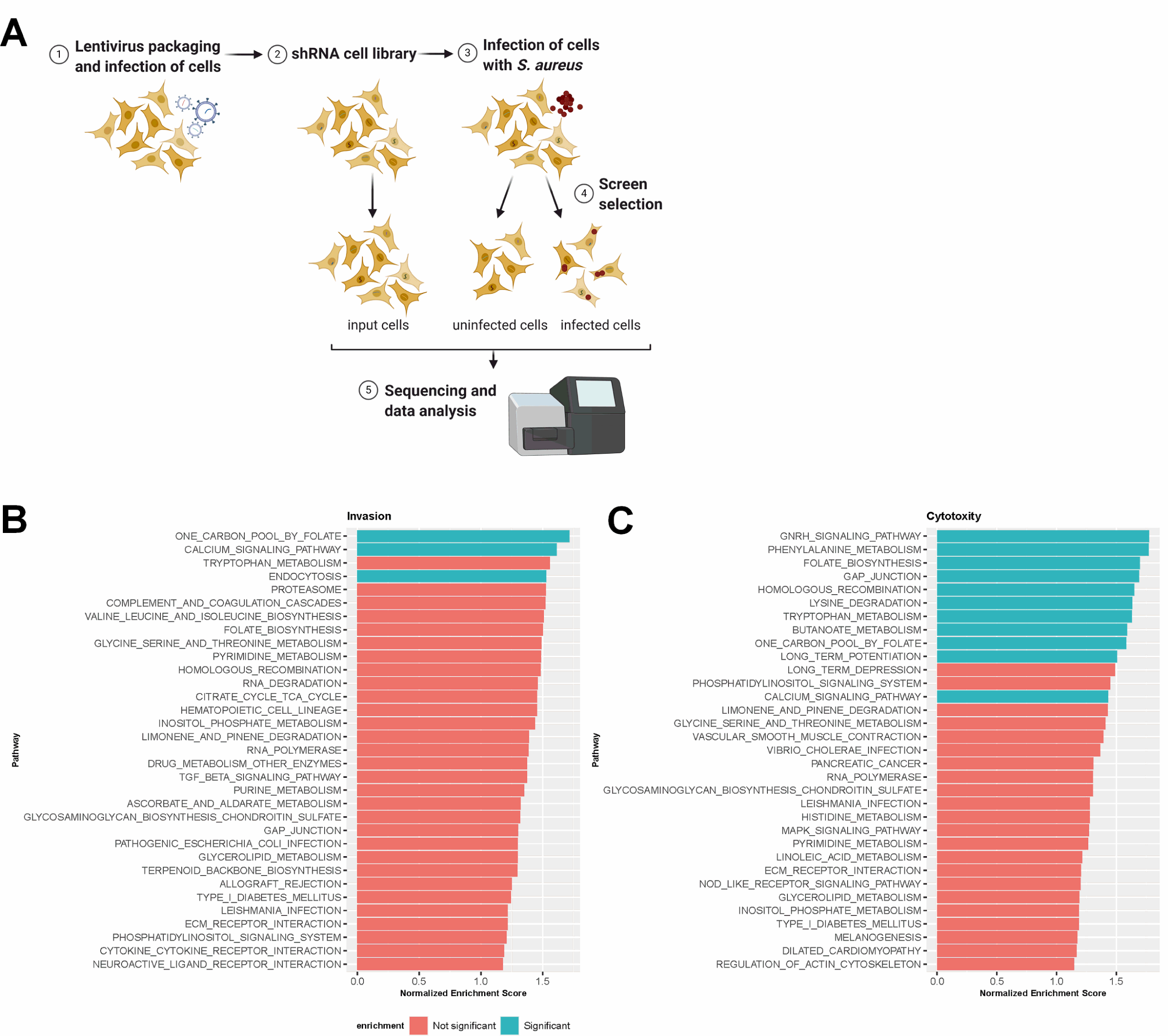
shRNA screen to identify host cell genes involved in *S. aureus* invasion and intracellular cytotoxicity. (A) Layout of shRNA screening approach. HeLa cells were first transduced with the shRNA library. Stable shRNA-HeLa cells were infected with *S. aureus* 6850 mRFP. 4.5 h p.i. dead cells were removed and uninfected and infected cells were sorted via FACS. Subsequently, genomic DNA was extracted and prepared for Illumina sequencing to determine enrichment or depletion of shRNAs in comparison to input HeLa cells, which were used as negative control. (B, C) A detailed Gene Set Enrichment Analysis (GSEA) of genes identified by the shRNA screen to be implicated in *S. aureus* invasion (B) or cytotoxicity (C) revealed key pathways. Pathways with *P*-value lower than 0.02 are noted as significant.

We next subjected the target genes of all differentially enriched or depleted shRNAs to gene set enrichment analysis (GSEA) (Figure 1B and C, Table S2). Among the pathways significantly enriched in the uninfected cell population, we identified the endocytosis pathway. Other pathways required for *S. aureus* cytotoxicity were Gonadotropin-releasing hormone (GNRH) signaling, phenylalanine metabolism, genes related to GAP junctions, homologous recombination, lysine degradation, tryptophan metabolism, butanoate metabolism and long-term potentiation. We identified two pathways, the one carbon pool by folate/folate biosynthesis as well as the calcium signaling pathway, where shRNAs were enriched in both *S. aureus* invasion and cytotoxicity.

### Intracellular *S. aureus* induce increase in cytoplasmic Ca^2+^

The GSEA analysis revealed the calcium signaling pathway to be involved in both *S. aureus* invasion and cytotoxicity. In addition, pathways connected to calcium signaling like long-term potentiation, implying ER calcium-channels and Ca^2+^-sensing and -regulated proteins, GNRH signaling, which is known to increase intracellular Ca^2+^ levels and PKC activation (47), and gap junctions, intercellular connection permeable for e. g. calcium ions, were enriched for the *S. aureus* intracellular cytotoxicity gene set. Therefore, we investigated, if intracellular *S. aureus* interfered with host cell Ca^2+^ signaling and thus may activate signaling pathways and Ca^2+^- sensing proteins implicated in infection-induced host cell death. We therefore used the genetically encoded fluorescent calcium sensor R-Geco (48), and generated a transgenic HeLa cell line in order to monitor changes in cellular Ca^2+^ concentration after infection with *S. aureus* by time-lapse microscopy. Extracellular bacteria were removed by treatment with lysostaphin. We observed a massive increase of the cytosolic Ca^2+^ concentration after the onset of intracellular bacterial replication and host cell shrinkage, which was followed by the formation of membrane blebs and a subsequent loss of bacterial and cellular fluorescence (Figure 2A, Video S1). Relative quantification of the cytosolic Ca^2+^ concentration demonstrated a roughly two- to six-fold increment of R-Geco fluorescence in infected cells between 5 and 7 h p.i. (Figure 2B). The high cytosolic Ca^2+^ level persisted for several minutes and up to one hour. No major changes in cellular Ca^2+^ concentrations were detected in uninfected cells (Figure 2B, Video S1). This was independent of the used bacterial strains, since we observed that the methicillin-resistant and cytotoxic strain JE2 induced a cytoplasmic Ca^2+^-rise, too (Figure 3F and G).

**Figure 2.**
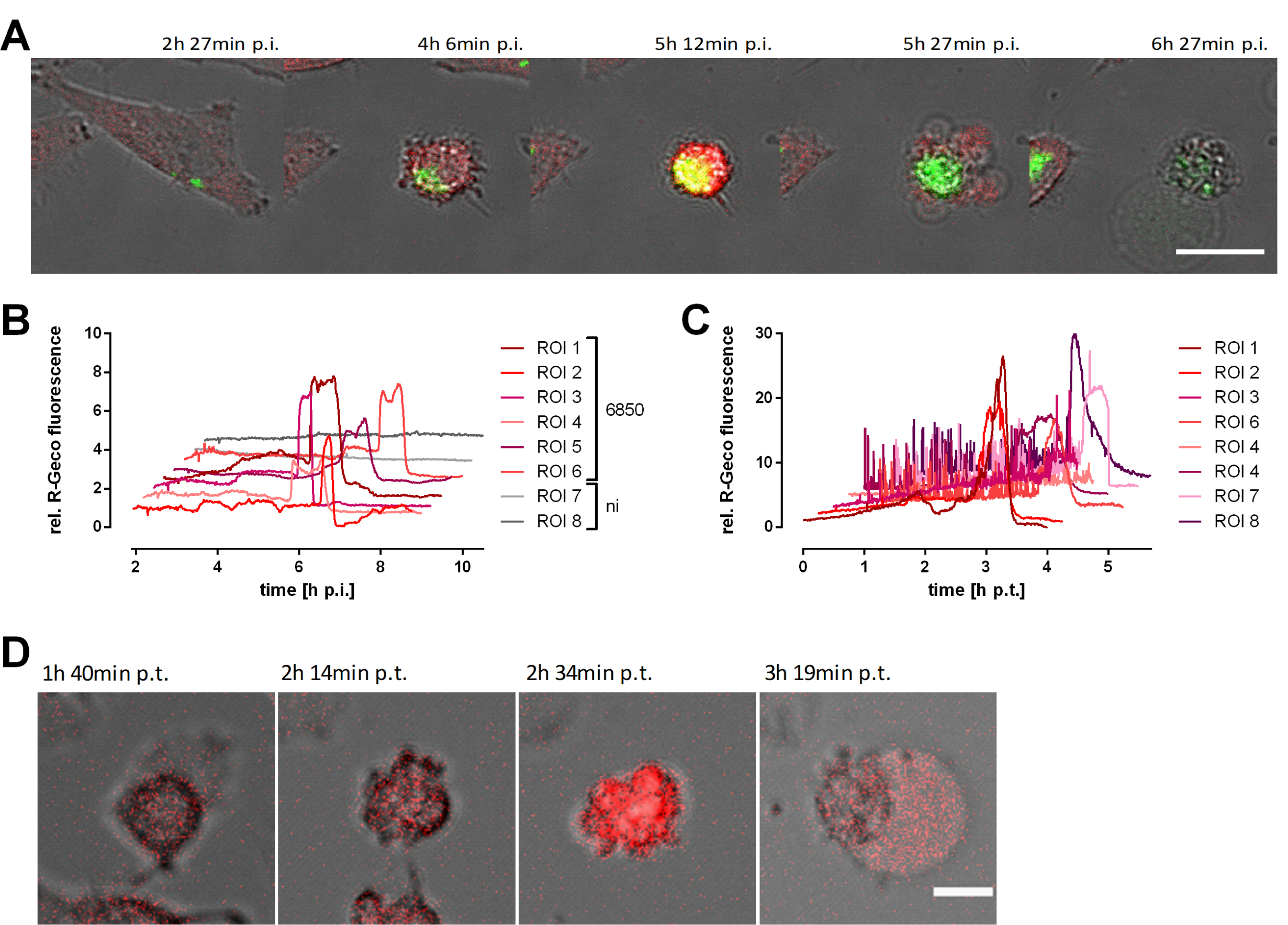
Infection with *S. aureus* or treatment with secreted *S. aureus* virulence factors leads to a massive cytosolic Ca^2+^-increase in epithelial cells. HeLa R-Geco cells were either infected with *S. aureus* 6850 GFP (A, B) or treated with 10 % sterilized supernatant of a *S. aureus* 6850 overnight culture (C, D) and visualized by live cell imaging. (A) Stills from time-lapse imaging are shown (green: *S. aureus*, red: R-Geco, gray: brightfield, scale: 25 µm). (B) Quantification of relative R-Geco fluorescence of single cells (ROI) infected with *S. aureus* 6850 or not infected (ni) was performed over the course of infection. (C) Relative R-Geco fluorescence of single cells was quantified upon SNT treatment. (D) Representative stills of one SNT-treated HeLa R-Geco cell (red: R-Geco, gray: brightfield, scale bar: 10 μm).

**Figure 3.**
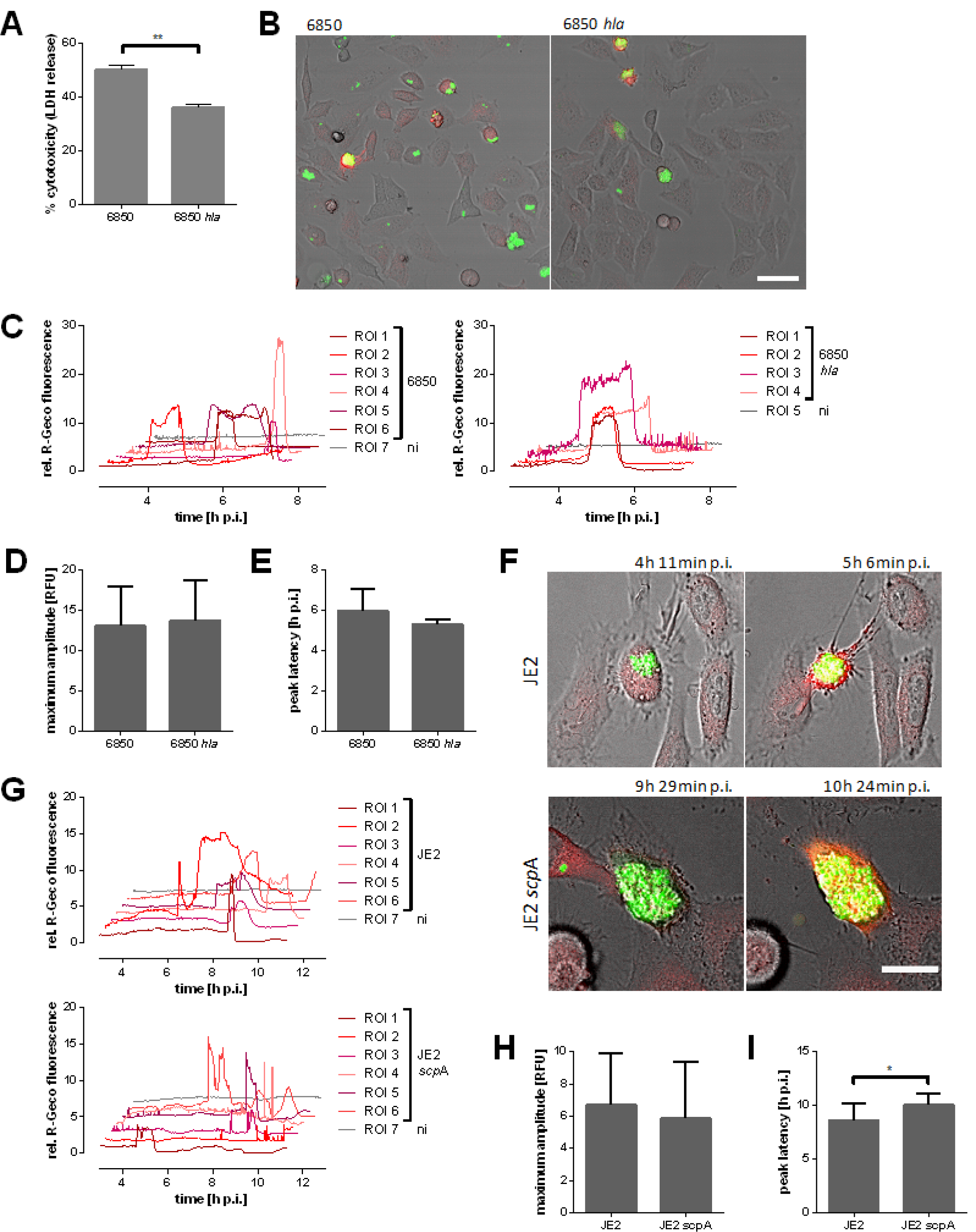
Ca^2+^ signaling in epithelial cells infected with *S. aureus* 6850 *hla* or JE2 *scp*A. (A) HeLa cells were infected with *S. aureus* 6850 or 6850 *hla* and cytotoxicity was determined at 6 h p.i. by LDH release. (B-I) HeLa R-Geco cells were infected with *S. aureus* 6850 GFP, 6850 *hla* GFP, JE2 GFP or JE2 *scp*A GFP and cytosolic Ca^2+^ concentration, i.e. R-Geco fluorescence, was measured by time-lapse imaging. (B) Stills of time-lapse imaging at 5 h 6 min p.i. are shown (green: *S. aureus*, red: R-Geco, gray: brightfield, scale: 50 µm). (C) Relative R-Geco fluorescence was quantified over the time period of infection in single uninfected cells (ni) or cells infected with *S. aureus* 6850 or *S. aureus* 6850 *hla*. (D) The peak amplitude of the relative R-Geco fluorescence of 4 to 12 single infected cells was determined. (E) The latency of relative R-Geco fluorescence peak after *S. aureus* intracellular infection was calculated in 4 to 12 single cells. (F) Representative stills from time-lapse fluorescence microscopy (green: *S. aureus*, red: R-Geco, gray: brightfield, scale bar: 20 µm). (G) Relative quantification of cytosolic Ca^2+^ concentrations of single infected (JE2) or uninfected (ni) cells. (H) The maximum amplitude of relative R-Geco fluorescence of 10 single infected cells was determined. (I) The latency after infection until the maximum amplitude of relative R-Geco fluorescence was quantified in 10 single cells. Statistical significance was determined by unpaired t-test (\**P*<0.05).

Since several *S. aureus* toxins were shown to induce Ca^2+^ flux in human cells (36), we treated HeLa R-Geco cells with 10 % sterile-filtered *S. aureus* culture supernatant. This led to high and frequent oscillations of cytosolic Ca^2+^ in most cells, which was followed by a sustained Ca^2+^ increase (Figure 2C, Video S2). Similar to intracellular infection with *S. aureus*, this cytoplasmic Ca^2+^ overload occurred after cell contraction, but prior to the formation of membrane blebs (Figure 2D) supporting that a secreted virulence factor may trigger Ca^2+^ fluxes from within the host cell.

The *S. aureus* pore-forming toxin alpha-hemolysin (*hla*) is known to form Ca^2+^- permissive pores and thus cytosolic Ca^2+^ elevations in different types of host cells (49, 50). Additionally, alpha-toxin plays a role in intracellular cytotoxicity of *S. aureus* (Figure 3A) (27, 51). Therefore, we investigated the Ca^2+^-flux inducing properties of an isogenic *S. aureus hla* deletion-mutant. Interestingly, infection of HeLa R-Geco cells with 6850 Δ*hla* did not abolish the cytosolic Ca^2+^ increase (Figure 3B and C). Further, the amplitude as well as the latency of maximal R-Geco fluorescence was not significantly different in HeLa cells infected with *S. aureus* Δ*hla* when compared to the wild type (Figure 3D and E). Another virulence factor, which we recently identified to play a role in *S. aureus* intracellular cytotoxicity, is the cysteine protease staphopain A (*scpA*) (31). Mutation of *scp*A in JE2 strain also did not impair cytoplasmic Ca^2+^-increase (Figure 3F-H), however, the onset of increase in cytosolic Ca^2+^ rise was delayed by 1.5 h on average (Figure 3I) and was shorter. Further, we observed Ca^2+^ oscillations (Figure 3G). This is in concordance with previous findings, in that knockout of *scpA* delays *S. aureus* intracellular cytotoxicity (31).

### Infection with intracellular *S. aureus* causes influx of extracellular Ca^2+^ in the host cell

We next studied the origin of the calcium ions. A preliminary test showed that a combination of Ca^2+^-withdrawal and chelation with BAPTA resulted in a strong reduction of R-GECO-fluorescence upon addition of the calcium ionophore ionomycin (Figure S2A). Thus, HeLa R-Geco cells were infected with *S. aureus* 6850 and the cell culture medium was replaced with Ca^2+^- containing or Ca^2+^-free medium (including BAPTA) after bacterial invasion and phagosomal escape (3 h p.i.). Subsequently, infected cells were monitored for changes in Ca^2+^ concentration (Figure 4A and B). When Ca^2+^ was omitted from the medium and additionally chelated with BAPTA, the relative R-Geco fluorescence showed a significantly reduced maximal amplitude (*p*=0.0004) in *S. aureus*-infected cells (Figure 4C), whereas no temporal changes of the cytosolic Ca^2+^ peak were noticed upon omission of extracellular Ca^2+^ alone (Figure 4D). Differences in cell morphology were not observed under the tested conditions.

**Figure 4.**
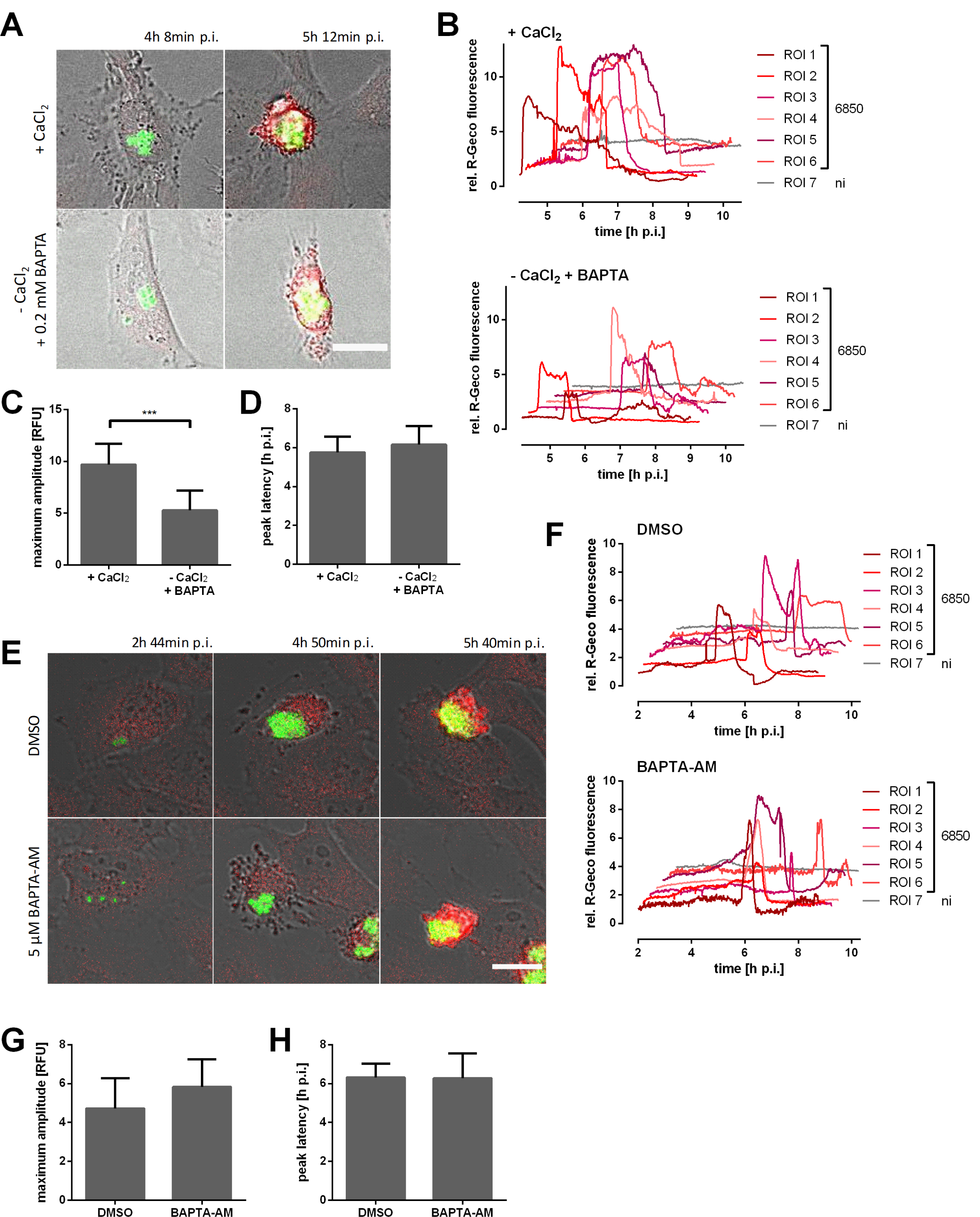
Removal of extracellular Ca^2+^ during *S. aureus* intracellular infection impairs cytosolic Ca^2+^-increase. HeLa R-Geco cells were infected with *S. aureus* 6850 GFP and 3 h p.i. medium with 1.8 mM CaCl_2_ (+ CaCl_2_) or without CaCl_2_ and 0.2 mM BAPTA (- CaCl_2_ + BAPTA) was added. Subsequently, R-Geco fluorescence was monitored by live cell imaging. (A) Representative stills of infected HeLa R-Geco cells with or without extracellular Ca^2+^ (green: *S. aureus*, red: R-Geco, gray: brightfield, scale bar: 20 µm). (B) Relative R-Geco fluorescence of single infected (6850) or uninfected (ni) cells was quantified over the time course of infection under the different conditions. (C) The maximal amplitude of the relative R-Geco fluorescence of 6 to 11 single infected cells under the different conditions was determined. (D) The latency of maximal relative R-Geco fluorescence peak after *S. aureus* intracellular infection was determined in 6 to 11 single infected cells. (E-H) HeLa cells were infected with *S. aureus* 6850 GFP, 2 h p.i. DMSO or 5 µM BAPTA-AM were added and R-Geco fluorescence was monitored over time. (E) Stills of representative infected cells are depicted (green: *S. aureus*, red: R-Geco, gray: brightfield, scale bar: 20 µm). (F) Relative R-Geco fluorescence of single infected (6850) or uninfected (ni) cells was quantified over the time course of infection under the different conditions. (G) The maximum amplitude of the relative R-Geco fluorescence of 6 to 11 single infected cells under the different conditions was determined. (H) The latency of relative R-Geco fluorescence peak after *S. aureus* intracellular infection in 6 to 11 single infected cells was calculated. Statistical analysis was performed by unpaired t-test (\*\*\**P*<0.001).

We next depleted intracellular Ca^2+^ with the cell-permeable chelator BAPTA-AM. Increasing concentrations of BAPTA-AM delayed, but did not reduce ionomycin-induced cytosolic Ca^2+^ increase (Figure S2B). Further, concentrations higher than 10 µM BAPTA-AM were cytotoxic to HeLa cells after several hours of incubation (data not shown). Therefore, we used 5 µM BAPTA-AM to treat HeLa R-Geco cells 2 h after *S. aureus* infection and R-Geco fluorescence was recorded over time. However, no differences in the cytoplasmic Ca^2+^ increase between BAPTA-AM treated and DMSO treated cells were observed (Figure 4E and F). Also, amplitude and latency of the maximal Ca^2+^ concentration were not significantly altered by BAPTA-AM treatment (Figure 4G and H). Thus, the Ca^2+^-chelating effect of BAPTA-AM was likely too small to interfere with *S. aureus*-induced cytosolic Ca^2+^ increase.

### Intracellular *S. aureus* perturbs ER Ca^2+^-homeostasis

Since Ca^2+^ release from intracellular stores of the host cell may also contribute to the cytoplasmic Ca^2+^ increase triggered by intracellular *S. aureus*, we generated a reporter cell line stably expressing a red-fluorescent endoplasmic reticulum (ER)- targeted fluorescent Ca^2+^ sensor (ER-LAR-Geco) (52) in addition to a green-fluorescent cytosolic Ca^2+^ indicator (G-Geco). The localization of ER-LAR-Geco was compartment specific with a typical distribution of the fluorophores to the ER (Figure S2C). We infected HeLa reporter cells with *S. aureus* 6850 and monitored fluorescence over time. In contrast to the cytoplasm, only small changes in ER Ca^2+^ concentration were observed in infected host cells (Figure 5A and B, Figure S2D). Quantification of ER-LAR-Geco fluorescence disclosed a decrease in ER Ca^2+^ concentration concomitant with the onset of cytosolic Ca^2+^ rise, whereas ER-LAR-Geco fluorescence increased minutes later to then drop again with the decrease of G-Geco fluorescence. When compared to the changes in cytosolic Ca^2+^ concentration, the shift of the ER-LAR Geco fluorescence was minor. To determine the dynamic range of the ER Ca^2+^ indicator, we treated HeLa ER-LAR-Geco G-Geco cells with Ca^2+^ perturbing agents. Ionomycin and thapsigargin also induced only small changes in ER-LAR-Geco fluorescence when compared to G-Geco (Figure S2E). These results show that the ER Ca^2+^ indicator was functional and that even high concentrations of Ca^2+^ perturbing agents only trigger a small increase in ER- LAR-Geco fluorescence.

**Figure 5.**
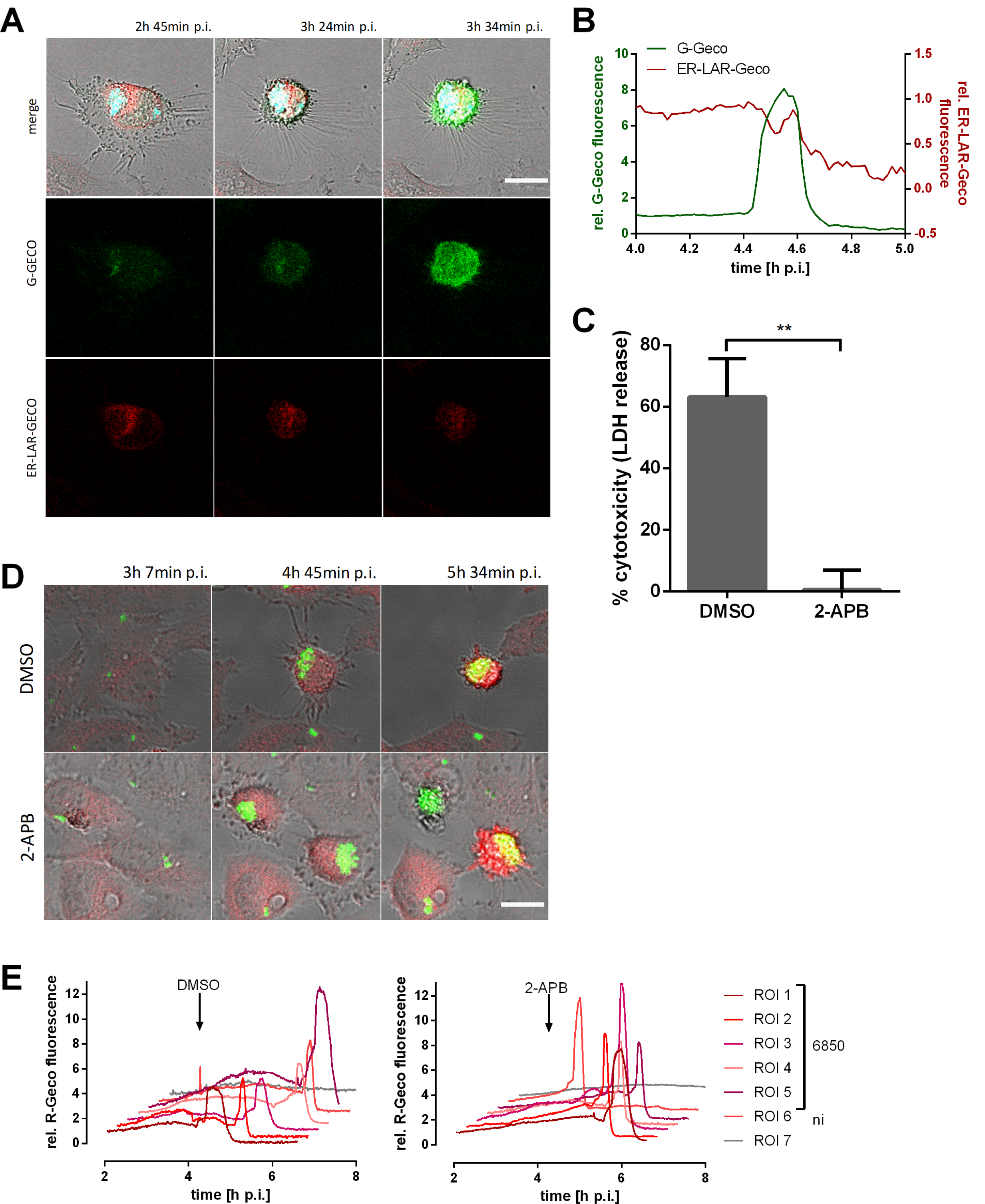
Intracellular *S. aureus* modulates ER Ca^2+^ concentration. (A, B) HeLa ER-LAR-Geco G-Geco cells were infected with *S. aureus* 6850 Cerulean and changes in ER and cytosolic Ca^2+^ concentration were monitored by live cell imaging. (A) Representative stills from time-lapse fluorescence microscopy (cyan: *S. aureus*, red: ER-LAR-Geco, green: G-Geco, gray: brightfield, scale bar: 20 µm). (B) Relative quantification of Ca^2+^ concentrations in cytosol and ER of a single infected cell. (C) HeLa cells were treated with 30 μM 2-APB or DMSO as solvent control 1 h prior to infection with *S. aureus* 6850 and cell death was determined at 6 h p.i. by LDH quantification. (D, E) HeLa R-Geco cells were infected with *S. aureus* 6850 GFP and cytosolic Ca^2+^, i.e. R-Geco fluorescence, was measured by time-lapse imaging. At 3 h 7 min p.i. DMSO or 30 µM 2-APB were added. (D) Representative stills from time-lapse fluorescence microscopy (green: *S. aureus*, red: R-Geco, gray: brightfield, scale bar: 20 µm). (E) Relative quantification of cytosolic Ca^2+^ concentrations of single infected (6850) or uninfected (ni) cells upon DMSO (left) or 2-APB (right) treatment.

Our observation of a simultaneous decrease in ER Ca^2+^ concentration and increase in cytosolic Ca^2+^ levels (Figure 5B, Figure S2D) thus suggest that store-operated calcium entry (SOCE) may be implicated in *S. aureus*-induced cell death. We therefore blocked SOCE with 2-aminoethyl diphenylborinate (2-APB). HeLa cells were treated with 30 µM 2-APB 1 h before infection and were subsequently infected with *S. aureus* 6850 in the presence of the inhibitor. Quantification of LDH release revealed that 2-APB treatment significantly (*p*=0.0015) reduced cytotoxicity of *S. aureus* by 99 % when compared to DMSO-treated cells (Figure 5C). This effect of 2-APB was reproducible in other epithelial cell lines, such as A549 and 16HBE14o- cells, infected with *S. aureus* 6850 (Figure S3A and B). Similarly, cytotoxicity of *S. aureus* JE2 in HeLa cells was abrogated by 2-APB treatment (Figure S3C).

Host cell invasion and phagosomal escape of *S. aureus* are prerequisites of intracellular cytotoxicity. We found that both were not significantly affected by 2-APB treatment (Figure S3D, E and F), although we noticed a small difference and non-significant in intracellular bacteria 3 h p.i. in 2-ABP-treated samples when compared to cells treated with the solvent control (Figure S3F). This difference was even more pronounced at 6 h p.i. (Figure S3G) suggesting that the bacteria were replicating less efficiently within the host cytosol. In cell culture medium, 30 µM 2-APB had only a mild inhibitory effect on *S. aureus* replication (Figure S3H).

Next, HeLa R-Geco were infected *S. aureus* and 2-APB was only added to infected cells 3 h p.i. to minimize effects on bacterial intracellular replication (Figure 5D and E). Time-lapse imaging showed that intracellular *S. aureus* was replicating despite 2- APB treatment (Figure 5D). However, addition of the inhibitor did not abolish *S. aureus*-triggered cytosolic Ca^2+^ increase (Figure 5E). Both amplitude and latency of the cytosolic Ca^2+^ increase were not significantly altered by 2-APB treatment under these conditions (Figure S3I and J).

### Intracellular *S. aureus* induces mitochondrial Ca^2+^-increase and cell lysis

The second major intracellular Ca^2+^ store in addition to the endoplasmic reticulum (ER) are the mitochondria. To investigate changes in mitochondrial Ca^2+^- concentration during *S. aureus* infection, we generated a dual Ca^2+^-reporter cell line stably expressing a red-fluorescent mitochondria-targeted fluorescent Ca^2+^ sensor (Mito-LAR-Geco) (52) in addition to a green-fluorescent cytosolic Ca^2+^ indicator (G-Geco) (Figure S4A) and we infected these cells with *S. aureus*. Within *S. aureus*- infected cells, we detected an increase in mitochondrial Ca^2+^ concentration, which ensued minutes after the increase in cytosolic Ca^2+^ (Figure 6A and B, Figure S4B). Interestingly, after the initial rise no or only a partial decrease of mitochondrial Ca^2+^ was detected in contrast to G-Geco fluorescence.

**Figure 6.**
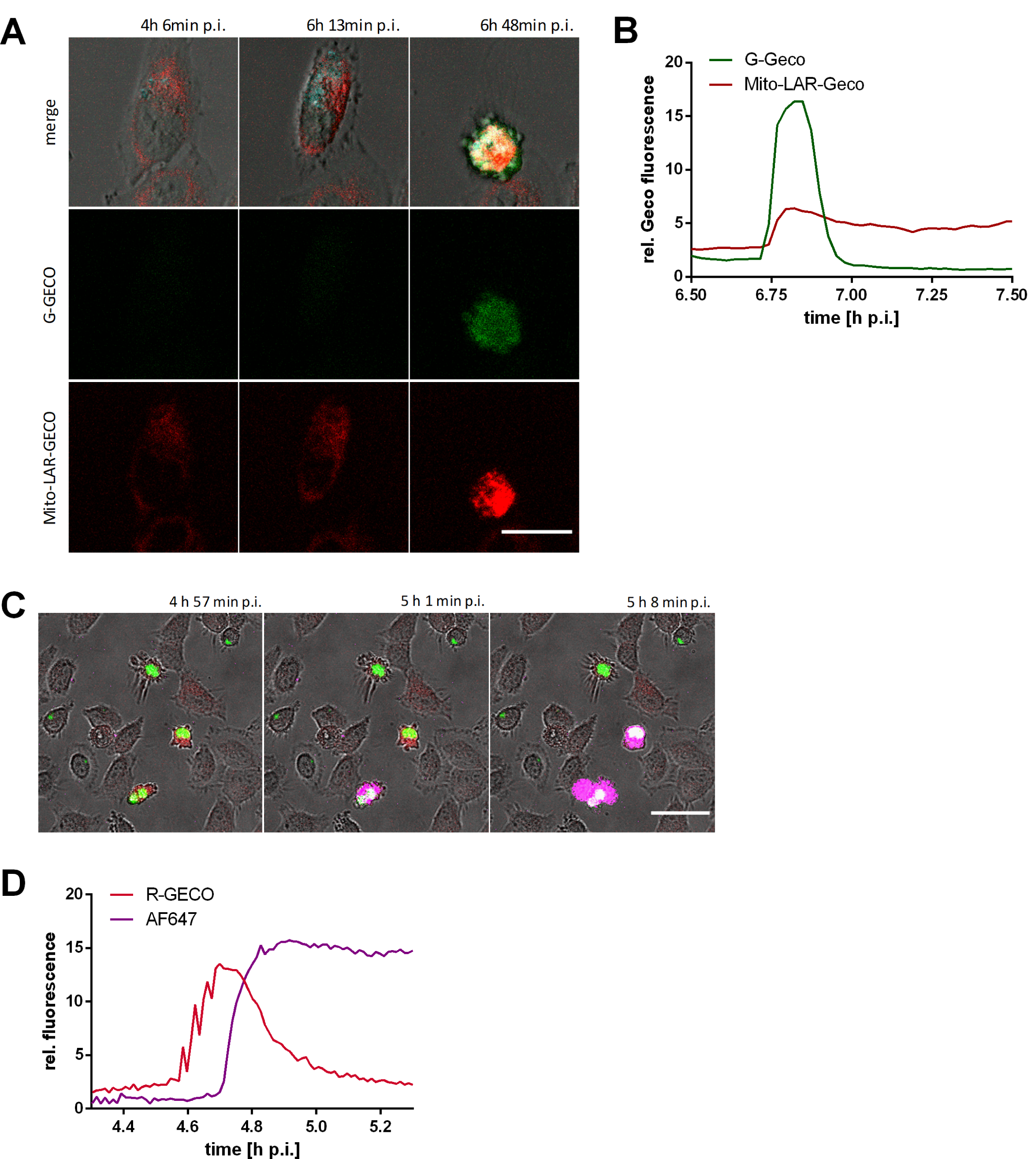
The *S. aureus* triggered cytosolic Ca^2+^ increase leads to a rise of mitochondrial Ca^2+^ concentration and cell lysis. (A, B) HeLa Mito-LAR-Geco G- Geco cells were infected with *S. aureus* 6850 Cerulean and changes in mitochondrial and cytosolic Ca^2+^ concentration were monitored by live cell imaging. (A) Representative stills from time-lapse fluorescence microscopy (cyan: *S. aureus*, red: Mito-LAR-Geco, green: G-Geco, gray: brightfield, scale bar: 25 µm). (B) Relative quantification of relative Ca^2+^ concentrations in cytosol and mitochondria of a single infected cell. (C, D) HeLa R-Geco cells were infected with *S. aureus* 6850 GFP and live cell imaging was performed after addition of the fluorescent dye AlexaFluor647 hydrazide. (C) Stills from time-lapse imaging (green: *S. aureus*, red: R-Geco, magenta: AF64, gray: brightfield, scale: 50 µm). (D) Relative fluorescence of a single cell infected with *S. aureus* 6850 was quantified over the course of infection.

Mitochondrial Ca^2+^-overload is associated with necrotic cell death. We therefore tested for plasma membrane permeability in *S. aureus*-infected HeLa R-Geco by addition of the small, membrane-impermeant fluorophore AlexaFluor647 hydrazide (AF647) to the culture medium. We observed that in infected cells influx of the fluorescent dye ensued after the increase in cytosolic Ca^2+^ (Figure 6C and S4C). Quantification of relative fluorescence of single infected cells revealed that AF647 fluorescence signal started to increase when R-Geco fluorescence was maximal levels (Figure 6D). By contrast, uninfected cells did not show evidence of AF647 influx.

### *S. aureus*-induced cytosolic Ca^2+^-rise is followed by caspase and calpain activation

*S. aureus*-induced cell death displays features of both, necrosis and apoptosis (53) and Ca^2+^ is also involved in regulation of apoptosis (54). Therefore, activation of effector caspases 3 and 7 was visualize by CellEvent™ Caspase-3/7 Green Detection Reagent (CellEvent Caspase 3/7) was added to HeLa R-Geco cells infected with *S. aureus* 6850. We observed that the increase in R-Geco fluorescence was followed by an increase in CellEvent Caspase 3/7 fluorescence (Figure 7A and B). Fluorescence quantification on a single cell level demonstrated that the effector caspases were activated in infected cells when the cytoplasmic Ca^2+^ signal peaked (Figure 7B and S4D). By contrast, fluorescence of CellEvent and R-Geco did not increase notably in uninfected cells (Figure S4D).

**Figure 7.**
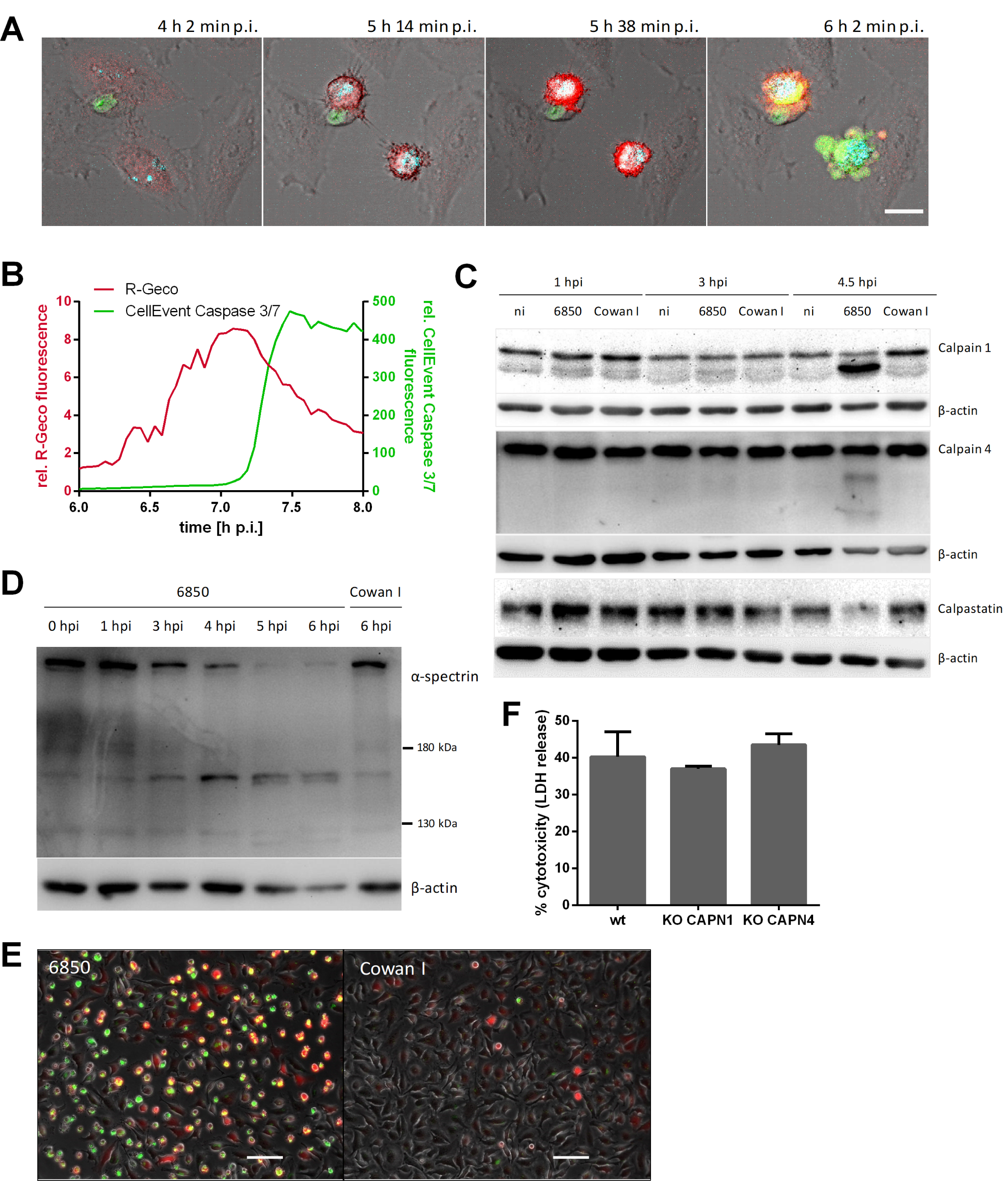
Activation of effector caspases and calpains in *S. aureus*-infected cells. (A, B) HeLa R-Geco cells were infected with *S. aureus* 6850 Cerulean and CellEvent™ Caspase-3/7 Green Detection Reagent (CellEvent Caspase 3/7) was added prior to live cell imaging. (A) Representative stills from time-lapse imaging are shown (cyan: *S. aureus*, red: R-Geco, green: CellEvent Caspase 3/7, gray: brightfield, scale bar: 20 µm). (B) The relative R-Geco and CellEvent Caspase 3/7 fluorescence of a single *S. aureus* 6850-infected cell was quantified over time. (C) HeLa cells were infected with *S. aureus* 6850 or non-cytotoxic Cowan 1 or remained uninfected (ni) and cell lysates were prepared at 1, 3 and 4.5 h p.i.. Proteins were separated by SDS-PAGE and transferred onto a PVDF membrane. Protein abundance and/or cleavage of calpain 1, calpain 4, calpastatin and β-actin as loading control was detected using the specific antibodies. (D) Immunoblot assay was performed as described for (C). *S. aureus* 6850- or Cowan 1-infected cells were lysed at 0, 1, 3, 4, 5 and 6 h p.i.. Calpain-specific cleavage of α-spectrin (150 and 145 kDa) was detected by application of an anti-α-spectrin antibody with anti-β-actin antibody as loading control. (E) HeLa cells were loaded with a fluorogenic calpain substrate (Boc-Leu-Met-CMAC) after infection with *S. aureus* 6850 or Cowan 1. The presence of activated calpains was detected at 4.5 h p.i. by fluorescence microscopy (red: Boc-Leu-Met-CMAC, green: *S. aureus*, gray: phase contrast, scale bar: 100 µm). (F) LDH release of HAP1 wild type (wt), CAPN4 KO or CAPN1 KO cells infected with *S. aureus* was determined at 6 h p.i. (n=2). Statistical analysis was performed by one-way ANOVA.

Calpains are Ca^2+^-activated cysteine proteases known to be implicated in cell death processes, such as apoptosis (42). Since calpain activation involves autocatalytic proteolysis (55) we investigated the degree of calpain 4 and calpain 1 proteolysis by Western blot 1, 3, and 4.5 hours after *S. aureus* infection (Figure 7C). An increase in autocatalytically processed calpains was not observed in uninfected cells or in cells infected with the non-cytotoxic control strain *S. aureus* Cowan 1, as well as in cell infected with cytotoxic *S. aureus* at early time points (1 and 3 h p.i.). At 4.5 h. p.i., by contrast, calpain autolysis products were detected. At this time, we also found reduced protein levels of the calpain-specific endogenous inhibitor calpastatin (Figure 7C). This suggested that calpains were activated in *S. aureus*-infected cells.

Since calpain-specific cleavage of the cytoskeletal protein α-spectrin is also used as a biomarker for calpain activation (56), we tested *S. aureus* 6850-infected HeLa cells by Western Blot for the calpain-specific breakdown products of α-spectrin (150 and 145 kDa). Protein bands with these molecular masses could be detected at 5 and 6 h p.i. (Figure 7D). By contrast, infection of HeLa cells with the non-cytotoxic Cowan 1 did not result in cleavage of α-spectrin. Additionally, a fluorogenic calpain substrate (Boc-Leu-Met-CMAC) specifically demonstrated calpain activity in host cells infected with cytotoxic 6850, but not with *S. aureus* Cowan 1 (Figure 7E).

We therefore deleted the genes for calpain 1 and calpain 4 (*CAPN1* and *CAPN4*, respectively) by a CRISPR/Cas9-based approach within the haploid HAP1 cell line, in which *S. aureus* demonstrates an intracellular pathogenicity comparable to that in HeLa cells (Figure S5A-E). As observed within HeLa, HAP1 cells demonstrated calpain activation as well as *S. aureus*-induced cytoplasmic Ca^2+^increase (Figure S5F-H). *CAPN1* and *CAPN4* gene knock-outs were verified by immunoblot (Figure S5I). However, the gene deletions did not reduce cytotoxicity of *S. aureus* (Figure 7F).

## Discussion

Previously conducted genome wide screens that investigated staphylococcal virulence factors mainly involved addition of purified toxins (57–59). However, *S. aureus* is readily internalized even by non-professional phagocytic host cells and is cytotoxic after phagosomal escape. We here employed a lentiviral shRNA library to identify host genes, which contribute to host cell death caused by the intracellular bacteria. Host cells, which were uninfected, were enriched for shRNAs targeting components of the endocytic machinery thereby supporting that *S. aureus* host cell invasion is a process driven by the host endocytosis machinery and thereby verifying the validity of our screening procedure.

Aside from calcium signaling pathway, “one carbon pool by folate” pathway and “folate biosynthesis” were identified to play are role in *S. aureus* invasion and intracellular cytotoxicity. Knock-down of factors involved in these pathways may result in metabolic and bioenergetic changes of the host cell and thus impair intracellular survival of *S. aureus*, which previously was shown to exploit the host cell carbon metabolism for intracellular replication (60, 61). Further, it was shown that the intracellular bacterium is able to alter the amino acid metabolism of its host cell (61). “Phenylalanine metabolism”, “tryptophan metabolism” and “lysine degradation” were identified by our screen to be crucial for cytotoxicity of intracellular *S. aureus*. Phenylalanine levels are increased in HeLa cells infected with *S. aureus* USA300 (60). Tryptophan metabolism is mediated by an enzyme called indoleamine 2,3- dioxygenase-1 (IDO1), which induces the conversion of tryptophan into kynurenine and downstream metabolites (62, 63). Kynurenine was shown to promote apoptosis of human and mouse neutrophils and inhibit production of ROS (64). Similarly, kynurenine may be utilized by intracellular *S. aureus* to induce cell death in epithelial cells. The specific relationship between these pathways and intracellular infection with *S. aureus* remains to be elucidated.

When comparing our results to a previously published shRNA screen of *S. aureus*- infected cells (65) genes involved in long term potentiation were identified in both studies. Inhibition of ER Ca^2+^-refill by the SERCA-inhibitor thapsigargin impaired bacterial internalization, but not *S. aureus* intracellular replication or cytotoxicity (65). Our study, which additionally differentiated between invasion- and cytotoxicity- relevant factors, demonstrated that intracellular *S. aureus* induced a perturbance of intracellular Ca^2+^-homeostasis in epithelial cells culminating in host cell death. So far, bacterial manipulation of the host cell Ca^2+^-signaling and homeostasis was only shown to be involved in disintegration of epithelial barriers, cytoskeletal rearrangements required for cell binding or internalization of the pathogen or control of innate defense cells (66).

Secreted *S. aureus* virulence factors induced strong cytosolic Ca^2+^-oscillations and elevations when added extracellularly to HeLa cells (Figure 2C). We observed that a cytosolic Ca^2+^-overload occurred after cell contraction but before the formation of plasma membrane blebs, which morphologically resembled that of intracellular *S. aureus* infection (compare Figure 2A and D). Thus, virulence factors secreted by intracellular *S. aureus* may trigger the Ca^2+^-overload when released into the cytoplasm of the host cell. Although the inner leaflet of the plasma membrane is composed of different proteins and phospholipids when compared to the outer leaflet, our results suggest that receptor-independent pore formation of *S. aureus* toxins may still occur from the inner side of the plasma membrane. Alpha-hemolysin, for instance, has been shown to nonspecifically integrate into membranes at high concentrations and thereby induces Ca^2+^-permissive pores (49, 50, 67–71). However, infection of epithelial cells with an alpha-toxin mutant did not significantly reduce or delay the cytosolic Ca^2+^ overload (Figure 3B-E) suggesting that other factors are responsible for the observed phenotype. Further, inactivation of the *S. aureus* cysteine protease staphopain A, which plays a role in *S. aureus* intracellular cytotoxicity (31), did not abolish cytosolic Ca^2+^ overload (Figure 3F-I).

We further observed that, in *S. aureus*-infected cells, ER Ca^2+^-levels decreased together with a cytosolic Ca^2+^-rise (Figure 5B and S5B) and that the latter also depended on calcium ions in the culture medium (Figure 4). This points to a role of store-operated Ca^2+^ entry (SOCE). During SOCE a release of Ca^2+^ from the ER triggers an influx of the ion from the extracellular space to refill the internal Ca^2+^ stores (41). In order to investigate the role of SOCE in the process of *S. aureus*- induced Ca^2+^ perturbations and host cell death, we applied 2-APB, which inhibits SOCE at higher concentrations (≥ 10-50 µM) (72, 73). Treatment of infected cells with 2-APB led to strong reduction of *S. aureus* intracellular cytotoxicity (Figure 5C), while it did not impair *S. aureus* invasion and phagosomal escape (Figure S6D-F). *S. aureus* intracellular growth was strongly attenuated upon inhibitor treatment (Figure S3F and G), which may imply that SOCE is not only required for induction of cytotoxicity but also for intracellular replication of *S. aureus*. 2-APB had only a minor direct effect on bacterial growth even over extended times (Figure S3H), which is in line with similar numbers of intracellular *S. aureus* in 2-ABP-treated cells (Figure S3F). When infected epithelial cells were treated with 2-APB only later during *S. aureus* intracellular infection (approx. 3 h p.i.), intracellular *S. aureus* was able to replicate (Figure 5D), but no effect of 2-APB on cytosolic Ca^2+^ increase was observed (Figure 5E and S3I and J). Therefore, no clear conclusion regarding the involvement of SOCE in *S. aureus*-induced cellular Ca^2+^ perturbations and the connection to host cell death can be drawn.

The rise in cytosolic Ca^2+^ was succeeded by increased Ca^2+^ levels in mitochondria (Figure 6B and S7B). Mitochondria are involved in Ca^2+^ compartmentalization and are able to buffer cytosolic Ca^2+^ to a certain extent. Upon high cytosolic Ca^2+^ concentrations larger amounts of Ca^2+^ can accumulate in the mitochondria (74) and excessive Ca^2+^ uptake by mitochondria may lead to mitochondrial permeability transition (MPT)-driven necrosis, which is characterized by abrupt loss of inner mitochondrial membrane osmotic homeostasis (75, 76). Cell lysis as a characteristic of a necrotic type of cell death was detected as a consequence of prolonged high cytosolic Ca^2+^ concentrations (Figure 6C and D). Accordingly, the cytosolic Ca^2+^ rise induced by intracellular *S. aureus* may lead to cell death by subsequent mitochondrial Ca^2+^ overload and MPT-driven necrosis.

Interestingly, it is speculated that the facultative intracellular pathogen *Shigella* induces mitochondrial Ca^2+^ overload by a sustained cytosolic Ca^2+^ increase due to plasma membrane permeabilization and thus MPT-driven necrotic cell death (77–79). However, so far only during host cell invasion of *Shigella* local and global Ca^2+^ fluxes were detected (80–82). Calpains, which are activated by Ca^2+^ release from intracellular stores likely act as executioner proteases during for MPT-driven necrosis. Additionally, in *S. aureus*-infected keratinocytes degradation of the endogenous calpain inhibitor calpastatin and reduced bacterial cytotoxicity upon calpain inhibitor treatment was observed (83).

We found that calpains were also activated in *S. aureus*-infected HeLa cells (Figure 7C-E). However, knock-out of calpains 1 and 4 did not abolish or reduce *S. aureus*- induced host cell death (Figure 7F). The high and sustained concentration of cytosolic Ca^2+^ induced by intracellular *S. aureus* may induce several cell death pathways in parallel by excess stimulation of Ca^2+^-sensitive targets, such as phospholipases, proteases and endonucleases in the cytosol (84–86). The Ca^2+^- induced damage may also trigger necrotic cell death without involving specific signaling. In this scenario, knockout of calpains cannot restrain host cell death.

Indications for an involvement of an apoptotic mode of cell death in *S. aureus*- infected cells were found, too. The activation of effector caspases 3/7 followed after the increase in cytosolic Ca^2+^ (Figure 7A and B). *S. aureus*-infected macrophages are also killed through a pathway characterized by membrane blebbing and activation of caspases 3/7 followed by cell lysis (87). Therefore also an apoptotic mode of cell death, which readily transits to a necrotic state, such as secondary necrosis, is likely, too (88).

In summary, our study shows that intracellular *S. aureus* induces Ca^2+^ perturbations in the host cell (Figure 8). Both plasma and ER membranes become permeable for calcium ions and induce a cytoplasmic and subsequently a mitochondrial Ca^2+^- overload, which results in breakdown of the plasma barrier function and cell death. Further, we demonstrate that *S. aureus* activated calpains 1/4 and effector caspases after translocation from the endocytic compartment to the host cytosol. Several cell death pathways thus are executed in parallel by activation of multiple Ca^2+^-sensitive proteins and we found evidence for MPT-driven necrosis as well as apoptotic pathways. However, the damage induced by strong perturbations of the cellular Ca^2+^- homeostasis and Ca^2+^-overload may not allow for reliable detection of specific signaling pathways and therefore may explain the confusing picture of host cell death pathways activated by intracellular *S. aureus* in the literature (53).

**Figure 8.**
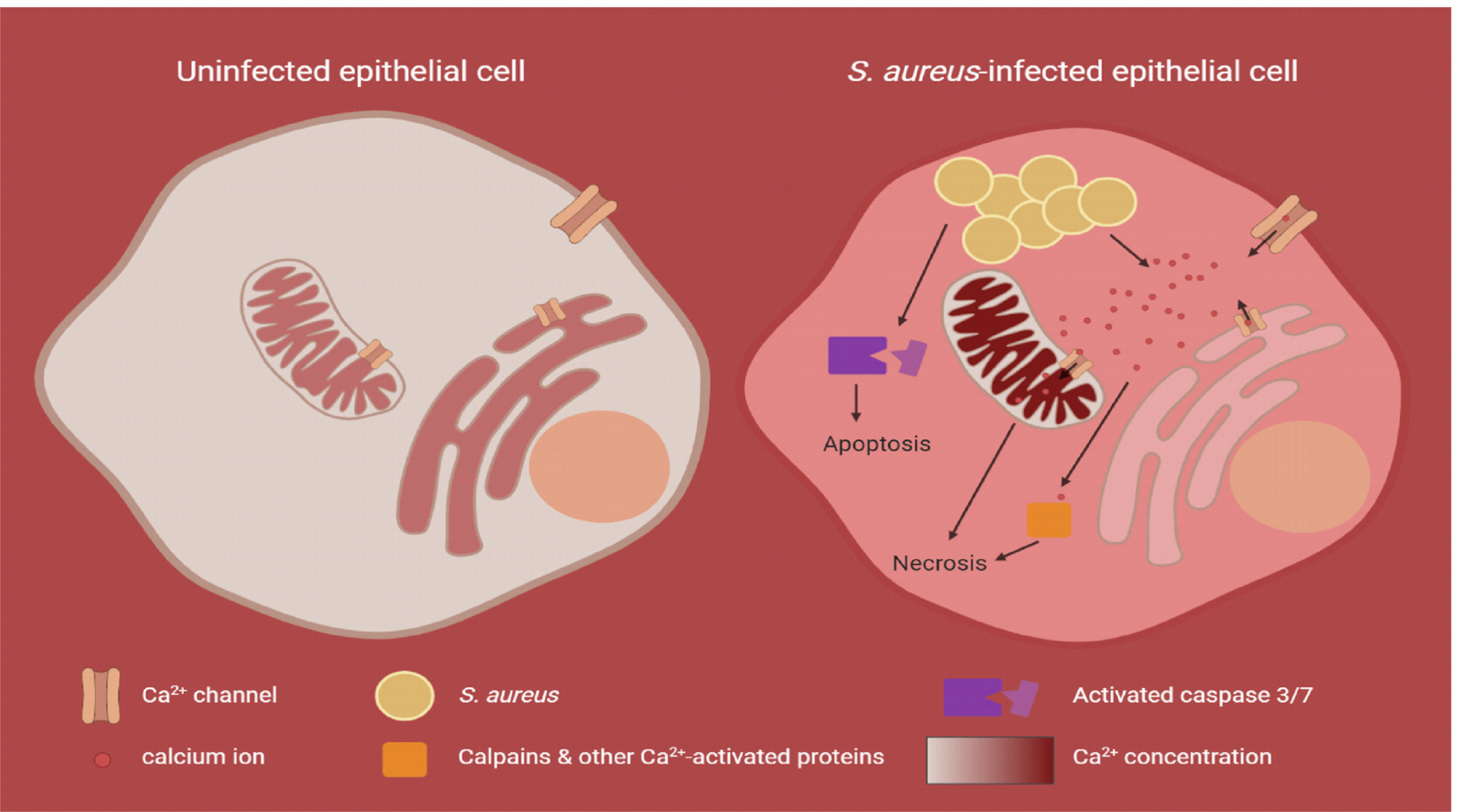
Graphical summary of *S. aureus*-induced perturbation of cellular Ca^2+^- homeostasis and subsequent host cell death. In uninfected epithelial cells, where the cellular Ca^2+^-homeostasis is not perturbed, the cytoplasmic Ca^2+^-concentration is 104-fold lower (approx. 100 nM) than the Ca^2+^-concentration in the extracellular milieu (approx. 1 mM). Intracellular organelles, like endoplasmic reticulum (ER) and mitochondria, known as the internal Ca^2+^-stores, accumulate Ca^2+^ and thus exhibit a higher Ca^2+^-concentration (100-500 µM) compared to the cytoplasm. Upon *S. aureus* infection, calcium ions from the extracellular space and ER are recruited to promote a cytoplasmic and subsequent mitochondrial calcium overload, which likely results in necrotic cell death. In parallel, excess stimulation of Ca^2+^-sensing proteins contributes to this effect. Additionally, effector caspases are activated by intracellular *S. aureus*, which execute apoptotic cell death.

## Methods

### Bacterial culture conditions

*Escherichia coli* strains were grown in Luria-Bertani broth (LB) and *Staphylococcus aureus* strains were grown in Tryptic soy broth (TSB, Sigma), if not stated otherwise. Media were supplemented with appropriate antibiotics, when necessary, and broth cultures were grown aerobically at 37°C overnight at 180 rpm. *E. coli* was selected on LB plates containing 100 µg/ml ampicillin and selective TSB plates for *S. aureus* were prepared using 10 µg/ml chloramphenicol and/or 10 µg/ml erythromycin.

For preparation of sterile supernatant, bacteria were grown overnight at 180 rpm in TSB and cultures were adjusted to an OD_600nm_ of 10. Subsequently, bacterial cultures were centrifuged, the supernatant was sterile filtered (0.22 µm pore size) and diluted to 10 % with cell culture medium.

Bacterial growth curves were measured using a TECAN plate reader. Bacterial cultures were inoculated in triplicates to an OD_600_ of 0.1 in 400 µl µl cell culture medium without FBS and grown for intended periods at 37 °C in a 48 well microtiter plate. Absorbance was recorded every 10 min at 600 nm.

### Generation of constructs

All used cell lines, strains, plasmids and oligonucleotides can be found in table S3 in the supplemental material.

In order to constitutively express the codon-adapted fluorescent reporter proteins mRFPmars and Cerulean we amplified the SarAP1 promoter from genomic *S. aureus* DNA by PCR using oligonucleotides SarAP1-f and SarAP1-r. The resulting PCR product was cloned into pCR2.1-Topo (Invitrogen), was sequence verified, restricted with SalI and KpnI, and eventually ligated into the plasmids pmRFPmars and pCerulean (89) opened with SalI and KpnI. We thereby exchanged the anhydrous tetracycline-inducible promoter including the ORF for the repressor TetR with the constitutively active SarAP1 promoter yielding pSarAP1-mRFP and pSarAP1- Cerulean, respectively. All assembled vectors were transformed into chemically competent *E. coli* DH5α and confirmed via PCR and Sanger sequencing (SeqLab, Göttingen). Subsequently, vectors were electroporated into *S. aureus* RN4220 and, if required, further either electroporated or transduced (using phage φ11) into the respective strains (Table S3).

For generation of stably integrating reporter gene expression vectors we used the lentiviral vector pLVTHM (90). Generation of the phagosomal escape reporter has been previously described (18, 91). Lentiviral vectors for expression of R-Geco, G-Geco, ER-LAR-Geco and Mito-LAR-Geco used following plasmids as PCR templates: CMV-R-GECO1, CMV-G-GECO1.1, CMV-ER-LAR-GECO1 and CMV-mito-LAR-GECO1.2, respectively (Table S3). The reporter genes were PCR amplified with oligonucleotides PmeI-pCMV-pLV and SpeI-BGHpolyA-pLV thereby introducing PmeI and SpeI restriction sites. PCR products were hydrolyzed with the respective restriction enzymes (Fermentas) and were ligated into correspondingly opened pLVTHM vector (90). Thereby replacing the original eGFP open reading frame with the respective sensor proteins. 20 µg of a plasmid preparation of these pLVTHM- derived plasmids were used in calcium phosphate-based co-transformation of a 15 cm dish of 293T cells along with 10 µg psPAX and 10 µg pVSVG. DMEM growth medium was exchanged after 4–8 hours. Two days after transfection, the supernatant was harvested and sterile-filtered (0.45 µm filter). HeLa and HAP1 cell lines were transfected with the virus particle preparations in presence of 10 µg/ml polybrene and after three sub-culturings the resulting transgenic cell lines were sorted on a FACSAria III cell sorter (BD).

The *hla* deletion mutant of strain 6850 was constructed using allelic replacement using the plasmid pKOR1 (92). A gene replacement cassette encompassing genomic regions upstream and downstream of the *hla* ORF was amplified by PCR with oligonucleotides attB1-hla-up-f and hla-up-r and hla-down-f and hla-down-r, respectively. Purified PCR products were restricted with SacII, ligated and subsequently recombined into pKOR1 using BP clonase (Invitrogen) and were subsequently transformed into competent *E. coli*. The resulting vector, pKOR1-Δ*hla*, was electroporated in *S. aureus* RN4220 (93), which accepts foreign DNA, and eventually in the target strain 6850. Selection of mutants was performed as described by T. Bae and O. Schneewind (92) and mutants were screened for successful gene replacement by PCR with the oligonucleotides hla-test-f and hla-test-r. Mutants were further checked for the phenotypic loss of β-hemolysis on sheep blood agar.

### shRNA screen

For generation of the HeLa-shRNA library 293T cells were cultivated in five 15 cm dishes to a density of 80 %. Every dish was transfected with 10 μg of pGIPZ lentiviral shRNAmir Library encoding for 10,000 different shRNAs, 7 μg of psPAX2 and 3.5 μg of pMD2.G using polyethylenimine (1 μl/1 μg of total DNA). Virus containing supernatant was collected 48 h and 72 h post transduction, sterile filtered and 1 μg/μl polybrene was added. HeLa cells cultivated in five 15 cm dishes were infected with the produced virus at a MOI of 0.1. 24 h post infection medium was changed and cells were cultivated with puromycin containing medium for selection. These five HeLa-shRNA sub-libraries encoding each for 10,000 different shRNAs were cultivated under puromycin pressure until they were used for infection.

One day before infection 3×10^6^ HeLa-shRNA cells of each of the five sub-groups were seeded in seven 15 cm dishes. Six dishes were infected with *S. aureus* 6850 mRFP at a MOI of 30 and 20 µg/ml lysostaphin was added 1 h p.i. in order to kill extracellular bacteria. 4.5 h p.i. dead cells were removed by two PBS washing steps and adherent viable cells were collected for FACS. Viable cells were selected via forward and sideward scatter (FSC-A and SSC-A) (Figure S1A). HeLa-shRNA cells, which express GFP, were sorted for the mRFP-positive (infection with *S. aureus* 6850) and mRFP-negative (uninfected) population using a FACS Aria III. GFP fluorescence was measured using a 488 nm laser and a 530/30 nm band pass filter for detection and mRFP fluorescence using 561 nm laser and a 610/20 nm band pass filter. One 15 cm dish of uninfected cells served as input control.

Genomic DNA of the sorted uninfected and infected cells and input cells was isolated with the Multisource Genomic DNA Miniprep Kit (Axygen Biosciences). The shRNA cassettes were amplified via PCR (primer see Table S3) and separation from the genomic DNA was obtained via agarose gel electrophoresis. Samples were diluted to a final concentration of 2 ng/μl and experion analysis confirmed the purity and concentration of the DNA. Samples were prepared for Illumina sequencing as suggested by the company.

High throughput sequencing has been performed with Illumina GAIIx genome analyzer. FASTQ files were aligned with Bowtie1 to a reference library containing the expected 10,000 shRNA sequences of each sub-library. Alignment of the sequencing reads to the reference library was quantified. Gene features without hits in any of the libraries were eliminated prior to normalization in order to avoid bias. Data quality control and normalization were implemented in R. Batch effect among different libraries was evident by the RLE plot and carefully removed by RUV methods (94, 95) Further normalization and differential expression analysis were conducted with edgeR (96, 97) whereby the TMM method was applied.

To expose in more detail the systematic impact of the different pathways, we further conducted a gene-set enrichment analysis (98) according to the pathways of KEGG and hallmark (gene set obtained from MSigDB ver. 7.0 (99)). We ranked all the genes with a score defined by “Score = -log10 (P-value) * log2FC”. The enrichment was computed using a permutation number of 5000. The pathways with high positive enrichment score together with *P*-value lower than 0.02 have been carefully investigated.

### Infection of epithelial cells

HeLa, A549 and 16HBE14o^-^ and were grown in RPMI1640 medium (#72400021, ThermoFisher Scientific) and HAP1 cells were grown in IMDM medium (#21980032, ThermoFisher Scientific) each supplemented with 10 % FBS (Sigma Aldrich) and 1 mM sodium pyruvate (ThermoFisher Scientific) at 37 °C and 5 % CO_2_. For infection 0.8 to 1 × 10^5^ cells were seeded into 12 well microtiter plates 24 hours prior to infection. 1 hour prior to infection medium was renewed and, if required, treatment with 30 µM 2-APB (Sigma Aldrich) was applied.

Bacterial overnight cultures were diluted to an OD_600nm_ of 0.4 and incubated for 1 hour at 37 °C and 180 rpm to reach exponential growth phase. Then, bacteria were washed twice by centrifugation and used to infect HeLa cells at a multiplicity of infection (MOI) of 50, if not stated otherwise. After one hour co-cultivation extracellular bacteria were removed by 30 minutes treatment with 20 µg/ml lysostaphin (AMBI) followed by washing and further incubation in medium containing 2 µg/ml lysostaphin and, if required, 30 µM 2-APB until the end of experiment.

### Live-cell imaging

Phase contrast and fluorescence microscopic images of live cells were acquired with a LEICA DMR microscope connected to a SPOT camera using a 10x (Leica HC PL FLUOTAR, NA=0.32) or 20x (Leica C Plan, NA=0.3) objective and VisiView^®^ software.

For time-lapse imaging, HeLa cells were seeded in 8 well chamber µ-slides (ibidi) 24 hours prior to infection. Infection with fluorescent protein-expressing bacterial strains was performed at a MOI of 5, as described above. Time-lapse imaging of the samples was performed on a Leica TCS SP5 confocal microscope using a 20x objective (Leica HC PL APO, NA=0.7) or 40x (Leica HC PL APO, NA=1.3) or 63x (Leica HCX PL APO, NA=1.3-0.6) oil immersion objectives. Prior to imaging cell culture medium was substituted with imaging medium (RPMI1640 without phenol red containing 10% FBS and 2 µg/ml lysostaphin) containing inhibitors, fluorogenic enzyme substrates and/or chemicals, if indicated. For Ca^2+^-depleted conditions DMEM without calcium (#21068028, ThermoFisher Scientific) substituted with 0.2 mM BAPTA (Merck Millipore) was used. As positive control 1.8 mM CaCl_2_ were added to this medium. Chelation of intracellular calcium ions was performed using BAPTA-AM (Merck Millipore). 5 µM CellEvent™ Caspase-3/7 Green Detection Reagent (ThermoFisher Scientific) were applied for monitoring effector caspase activation and 2.5 µM AlexaFluor647 hydrazide for detecting cell lysis.

The µ-slides were transferred to a pre-warmed live-cell incubation chamber surrounding the confocal microscope and a temperature of 37 °C were applied during imaging. LAS AF software was used for setting adjustment and image acquisition. All images were acquired at a resolution of 1024×1024 pixels and recorded in 8-bit mode at predefined time intervals. In certain cases, chemical treatment was performed during imaging *in situ* at the live-cell incubation chamber. All image-processing steps were performed using Fiji (100). For quantification of fluorescence intensities raw imaging data was used. For single cell analysis one region of interest (ROI) was defined for all recorded time frames. Fluorescence intensities (mean of RFU) were measured, background was substracted and data were normalized to time point zero (R_0_) obtaining relative fluorescence values. Maximum amplitude was determined as highest relative fluorescence value measured and peak latency was defined as the time point of maximum amplitude.

### Cytotoxicity assays

HeLa cells were infected as described above. At the desired time point after infection medium, which possibly contained detached dead cells, was collected from the wells and adherent cells were detached using TrypLE (ThermoFisher Scientific). Adherent and suspension cells of each sample were pooled and after centrifugation for 5min at 800 x g cells were carefully resuspended in cell culture medium with 10 µl/ml 7-AAD (BD Biosciences). After 10 minutes incubation in the dark cells were immediately analyzed by flow cytometry using a FACS Aria III (BD Biosciences) and BD FACSDiva Software (BD Biosciences). Forward and sideward scatter (FSC-A and SSC-A) were used to identify the cell population and doublet discrimination was performed via FSC-H vs. FSC-W and SSC-H vs. SSC-W gating strategy. 7-AAD fluorescence was measured using a 561nm laser for excitation and 610/20 nm band pass filter for detection. 10,000 events were recorded for each sample. Uninfected cells served as negative control.

The LDH assay was performed as described previously (31). Briefly, HeLa cells were infected as described above and 1.5 hours after infection medium was replaced by RPMI1640 without phenol red containing 1 % FBS and 2 µg/ml lysostaphin. 6 hours after infection medium was removed from the wells, shortly centrifuged and 100 µl of supernatant of each sample were transferred into the well of a 96 well microtiter plate in triplicates. LDH release was measured using the Cytotoxicity Detection Kit Plus (Roche) according to manufacturer’s instruction.

### Invasion and intracellular replication assay

HeLa cells were infected with GFP expressing strains and prepared for flow cytometry as described above. Adherent and suspension cells were resuspended in fresh medium without phenol red containing 1 % FBS and 2 µg/ml lysostaphin. To determine invasion, infected cells at 1 h p.i. were treated with 20 µg/ml lysostaphin for 10 minutes to remove extracellular bacteria and the percentage of GFP-positive cells representing the infected cells was measured by flow cytometry. Intracellular replication was determined by measurement of GFP fluorescence (arbitrary units, AU) of the infected cells 1 and 3 hours after infection by flow cytometry using a FACS Aria III (BD Biosciences). The intensity of GFP fluorescence corresponds to the amount of intracellular bacteria. Gating and analysis were performed as described above. GFP fluorescence was measured using a 488nm laser and a 530/30 nm band pass filter for detection. Uninfected cells served as negative control.

### Phagosomal escape assay

Phagosomal escape was determined as described previously (31). HeLa YFP-cwt cells were infected with mRFP-expressing bacterial strains at a MOI of 10 in a 24 well µ-plate (ibidi). After synchronization of infection by centrifugation, infected cells were incubated for 1 hour to allow bacterial invasion. Subsequently, a 30 minute-treatment with 20 µg/ml lysostaphin removed extracellular bacteria, after which the cells were washed and medium with 2 µg/ml lysostaphin was added. Three hours after infection, cells were washed, fixed with 4 % paraformaldehyde overnight at 4 °C, permeabilized with 0.1% Triton X-100 and nuclei were stained with Hoechst 34580. Images were acquired with an Operetta automated microscopy system (Perkin-Elmer) and analyzed with the included Harmony Software. Co-localization of YFP-cwt and mRFP signals indicated phagosomal escape.

### Gene knock-out using CRISPR/Cas9

For designing gene-specific sgRNAs the CRISPOR online tool (crispor.tefor.net) was used and gene sequences were retrieved from Ensemble (www.ensembl.org). Exons coding for essential function(s) of the protein were chosen for genetic manipulation. Synthetized sgRNA oligonucleotides (Table S3) were cloned into pSp-Cas9(BB)-2A- GFP according to the protocol from F. A. Ran et al. (101). Insertion of the sgRNA was verified by Sanger sequencing (SeqLab, Göttingen) using primer U6-fwd.

For plasmid transfection, HAP1 cells with a low passage number were seeded into 6 well microtiter plates at a density of 5 × 10^5^ cells/well in IMDM medium (#21980032, ThermoFisher Scientific) containing 10 % FBS (Sigma Aldrich) and antibiotic 24 h before transfection. 3.6 µg pSp-Cas9(BB)-2A-GFP-sgRNA and 400 ng pSc-TIA- CMV-BSR-TIA were mixed with polyethylenimine and OptiMEM (#51985026, ThermoFisher Scientific) for transfection of cells. After 24 h cell were diluted and seeded in cell culture dishes (100 × 20 mm). The next day selection was applied by treatment with 30 µg/ml blasticidin. Subsequently, cells were monitored over 2- 3 weeks and, if necessary, cells were washed and medium was renewed. When cell colonies were large enough, they were transferred to a 12-well plate using trypsin- soaked, sterile Whatman paper. Gene knock-out was verified by western blot.

### SDS-PAGE and Immunoblotting

Protein samples from human cells were prepared in 2x Laemmli buffer (100 mM Tris/HCl (pH 6.8), 20 % glycerol, 4 % SDS, 1.5 % β-mercaptoethanol, 0.004 % bromophenol blue) and immediately incubated at 95°C for 10 min for protein denaturation. Proteins were separated via gel electrophoresis on 7.5-12 % polyacrylamide gel and transferred to a PVDF membrane (Sigma Aldrich) using a semi-dry blotting system. The PVDF membrane was incubated for 1 hour in blocking solution (5 % human serum in 1x TBS-T) and overnight at 4°C with the first antibody (diluted in blocking solution). Antibodies against Calpain 1 (1:1000, #2556, Cell signaling Technology), Calpain 4 (1:500, MAB3083, Merck Millipore), Calpastatin (1:1000, #4146, Cell signaling Technology), α-spectrin (1:300, sc-48382, Santa Cruz Biotechnology), β-actin (1:3000, A5441, Sigma Aldrich) and β-tubulin (1:1000, MAB3408, Merck Millipore) were used. Primary antibodies were detected with a horseradish peroxidase (HRP)-conjugated secondary antibody (1:3000, #170-6515, Biorad or #12-349, Merck Millipore, in 1x TBS-T with 5 % non-fat dry milk) using enhanced chemiluminescence (ECL) and an Intas imaging system (Intas Science Imaging).

### Calpain activity assay

HeLa cells were infected with *S. aureus* 6850 GFP as described above. At 4 h p.i. 10 µM CMAC peptidase substrate t-BOC-Leu-Met (ThermoFisher Scientific) was added to the cells and incubated for 30 min at 37 °C and 5 % CO_2_. Subsequently, phase contrast and fluorescence microscopic images of live cells were acquired with a LEICA DMR.

### Statistical analyses

Data were analyzed using GraphPad Prism Software (GraphPad Software, Version 6.01). For statistical analysis three biological replicates were performed, if not indicated otherwise. All data are presented as means with standard deviation (SD). P-values ≤0.05 were considered significant. Pairwise comparisons were assessed using unpaired Student’s t-test. Analysis of variance (ANOVA) was performed to determine whether a group of means was significantly different from each other. ANOVA was performed with Tukey’s post-hoc analysis for defining individual differences.

## Acknowledgements

We thank the German Research Foundation (DFG; http://www.dfg.de) for funding this project within the Transregional Research Center TRR34 under code C11 to M.J.F. and T.R. (K.S., A.C.W.). We thank the German Research Foundation (DFG; http://www.dfg.de) for funding T.D. (project number 374031971 - TRR240/Z2). We are further grateful to Kerstin Paprotka, Magdalena Grosz and Simone Vormittag for generation of constructs and to Ursula Eilers and the Core Unit Functional Genomics (University Würzburg) for support with Operetta Imaging. We thank Mark Onyango (University Gießen) for assistance in bioinformatical analysis. CMV-R-GECO1, CMV-G-GECO1.1, CMV-ER-LAR-GECO1, and CMV-mito-LAR-GECO1.2 were a gift from Robert Campbell (University of Alberta, Canada). We thank Lucas Jae (Gene Center, Munich) for the HAP1 cell line and plasmid pSc-TIA-CMV-BSR-TIA.

## Author Contributions

KS, ACW, LC, CPA, TD, MJF and TR conceived and designed the experiments. KS and ACW performed the experiments. KS and LC analyzed the data. KS, LC, TD, MJF and TR wrote the paper.

## Supporting information

**Table S1. Genes targeted by shRNAs differentially enriched in uninfected vs. input sample and in infected vs. input sample (P-value <0.001; Log2FC >1.5 or Log2FC < -1.5)**

**Table S2. GSEA of genes identified by the shRNA screen to be implicated in *S. aureus* invasion or cytotoxicity**

**Table S3.**
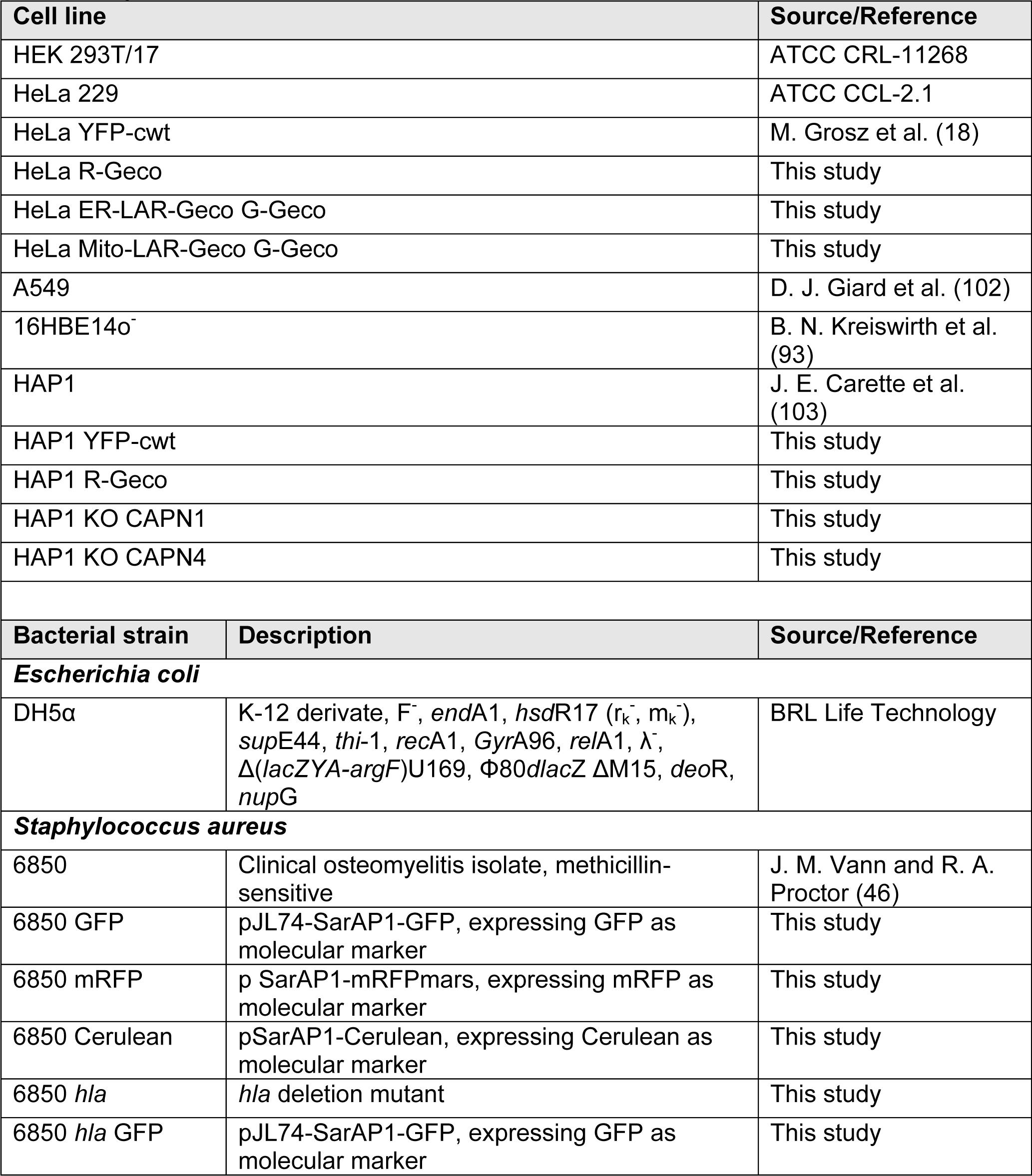
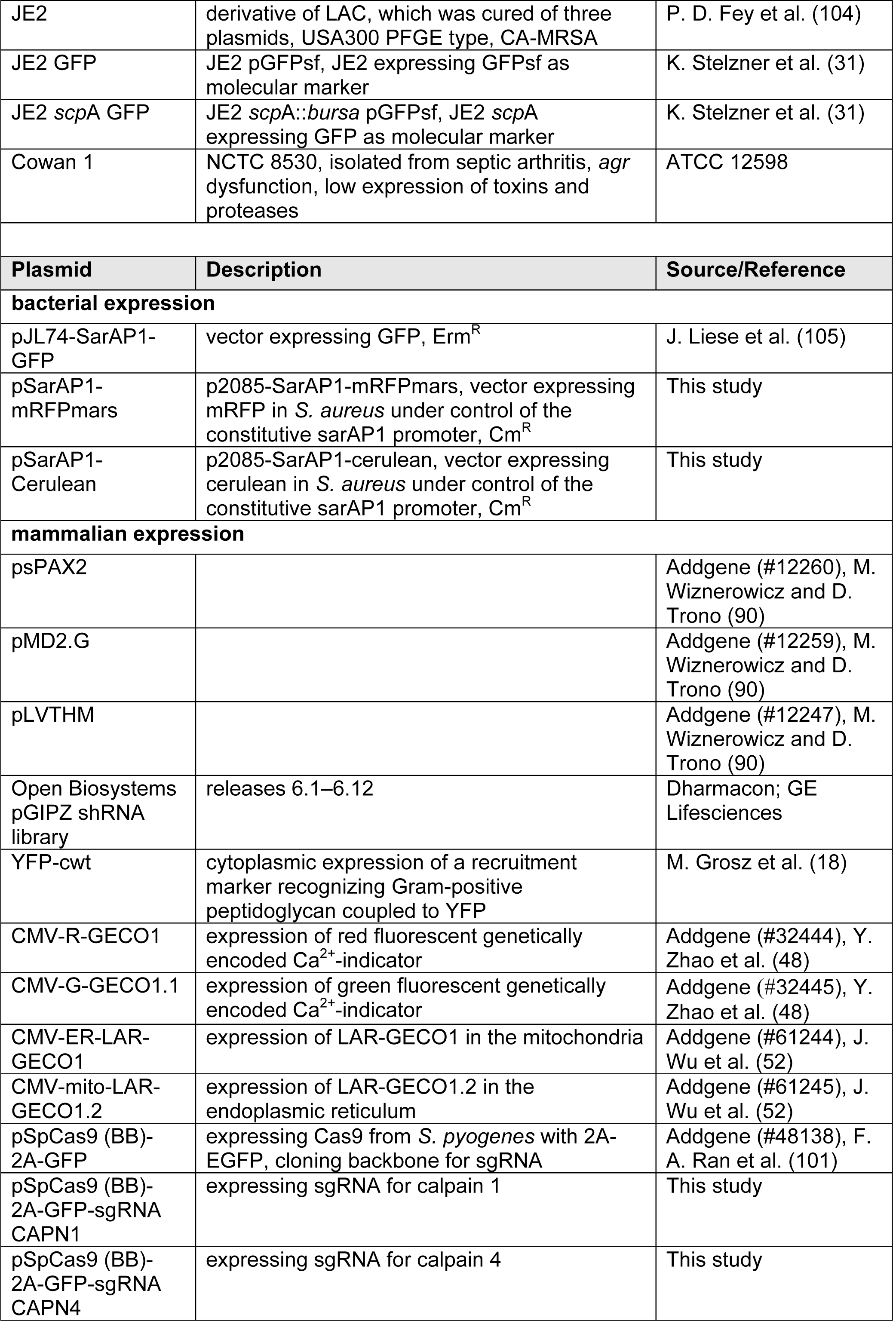
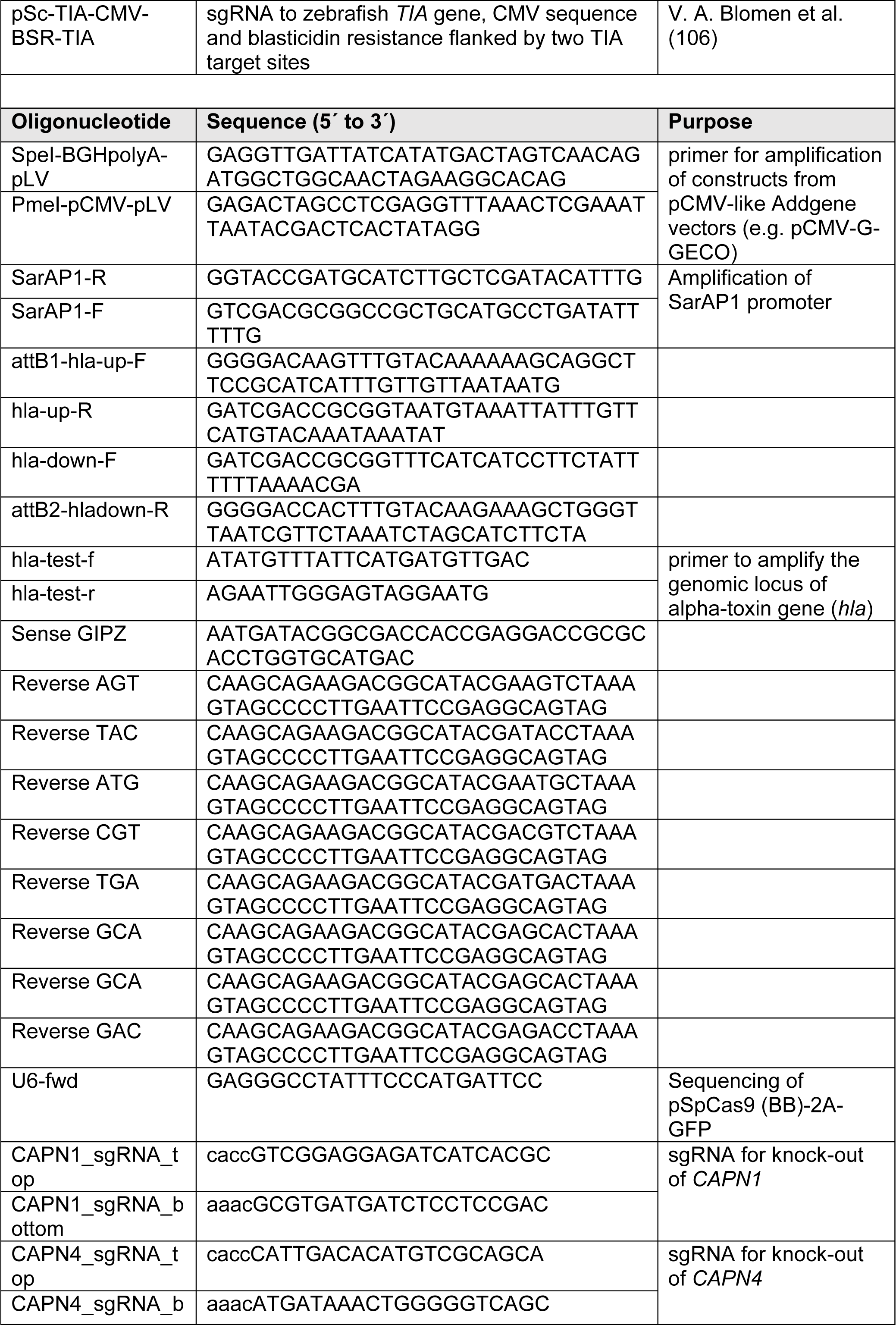
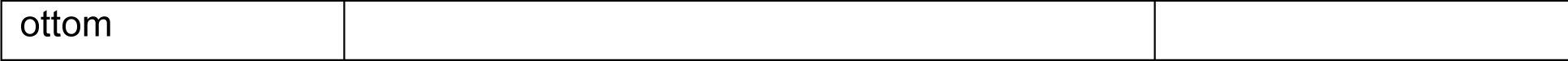
Cell lines, bacterial strains, plasmids and oligonucleotides used in this study.

### Supporting information captions

**Figure S1. shRNA screen**. (A) Intact cells were defined by flow cytometric-analysis via FSC and SSC (P1). GFP expressing cells, indicating shRNA expression, were gated as P4. Cells harboring intracellular *S. aureus* 6850 showed increased mRFP fluorescence. Uninfected and infected HeLa cells were gated as P5 and P6, respectively. (B, C) Quality control and normalization of shRNA libraries. Relative log expression plot examines the result of normalization method (C). Principal component analysis (PCA) plot of six shRNA libraries after removing batch effect. (D, E) Heatmaps of genes targeted by significantly enriched or depleted shRNAs when compared to the input library. (D) Comparison between uninfected and input library with a threshold of P-value 0.001 and Log2FC 3.5. (E) Comparison between the infected library and input library with a threshold of P-value 0.001 and Log2FC 3.

**Figure S2. Ca**^**2+**^**-flux via plasma and ER-membrane in *S. aureus*-induced cytoplasmic Ca**^**2+**^ **rise**. (A) HeLa R-Geco cells were cultivated in medium with 1.8 mM CaCl_2_, without CaCl_2_ or without CaCl_2_ and 0.2 mM BAPTA and 0.5 µM ionomycin were added while recording of R-Geco fluorescence. (B) HeLa R-Geco cells were treated with DMSO or 5, 10, 25 or 50 µM BAPTA-AM and while recording R-Geco fluorescence 0.5 µM ionomycin were added. (C) Fluorescence images of HeLa ER-LAR-Geco cell lines (BF: brightfield, scale bar: 20 µm). (D) HeLa ER-LAR-Geco G-Geco cells were infected with *S. aureus* 6850 Cerulean and changes in ER and cytosolic Ca^2+^ concentration were monitored by live cell imaging. Relative quantification of Ca^2+^ concentrations in cytosol and ER of single infected cells and an uninfected cell (lower right graph) are shown. (E) Fluorescence of HeLa ER-LAR- Geco G-Geco cells was recorded by live cell imaging, while 10 µM ionomycin (left) or 10 µM thapsigargin (right) were added (see arrows).

**Figure S3. 2-APB attenuates bacterial cytotoxicity, but not *S. aureus*-induced cytoplasmic Ca**^**2+**^**-increase**. (A-G) A549 (A), 16HBE14o^-^ (B), HeLa (C,D,F,G) or HeLa YFP-cwt (E) cells were treated with 30 µM 2-APB or DMSO 1 h prior to infection with *S. aureus* 6850 (A,B), JE2 (C), 6850 GFP (D, E, G) or 6850 mRFP (F). (A-C) LDH release was quantified 6h p.i.. (D) Number of infected host cells, i.e. GFP-positive cells, was measured at 1 h p.i.. (E) Translocation of bacteria into the host cell cytosol was determined by quantification of co-localization of intracellular bacteria and the cytosolic escape marker YFP-cwt at 4 h p.i. (n=2). (F) The relative number of intracellular bacteria was identified by measuring the mean fluorescence intensity of infected cells at 1 and 3 h p.i.. (G) Fluorescence microscopy images at 6 h p.i. (green: *S. aureus*, gray: phase contrast, scale bar: 100 µm). (H) Optical density (OD) at 600 nm of *S. aureus* 6850 treated with 30 µM 2-APB or DMSO in RPMI medium without FBS was determined for 17.5 h at 10 min-intervals. (I, J) HeLa R-Geco cells were infected with *S. aureus* 6850 GFP and R-Geco fluorescence was measured by time-lapse imaging. At 3 h 7 min p.i. DMSO or 30 µM 2-APB were added. (I) The peak amplitude of the relative R-Geco fluorescence of 7 to 11 single infected cells was quantified after DMSO or 30 µM 2-APB treatment. (J) The latency of the relative R-Geco fluorescence peak after *S. aureus* intracellular infection was determined and the mean value of 7 to 11 cells was calculated. Statistical analysis was performed by unpaired t-test.

**Figure S4. The cytosolic Ca**^**2+**^ **increase in *S. aureus*-infected cells is followed by mitochondrial Ca**^**2+**^**- increase, cell lysis and effector caspase activation**. (A) Fluorescence images of HeLa Mito-LAR-Geco G-Geco cell (BF: brightfield, scale bar: 20 µm). (B) HeLa Mito-LAR-Geco G-Geco cells were infected with *S. aureus* 6850 Cerulean and changes in mitochondrial and cytosolic Ca^2+^ concentration were monitored by live cell imaging. Relative Ca^2+^ concentrations in cytosol and mitochondria of single infected or uninfected (lower right graph) cells were quantified over time. (C) HeLa R-Geco cells were infected with *S. aureus* 6850 GFP and live cell imaging was performed after addition of the fluorescent dye AlexaFluor647 hydrazide. Relative fluorescence of single cells infected with *S. aureus* 6850 or single uninfected cells (lower right graph) was quantified over the course of infection. (D) HeLa R-Geco cells were infected with *S. aureus* 6850 Cerulean and CellEvent Caspase-3/7 was added prior to live cell imaging. The relative R-Geco and CellEvent Caspase 3/7 fluorescence of single uninfected (lower right graph) or *S. aureus* 6850- infected cells was quantified over time.

**Figure S5. The intracellular lifestyle of *S. aureus* in HeLa and HAP1 cells**. (A) HeLa and HAP1 cells were infected with *S. aureus* 6850 GFP and invasion was determined at 1 h p.i. by flow cytometry as percentage of infected (i.e. GFP-positive) cells. (B) For detection of phagosomal escape HeLa YFP-cwt and HAP1 YFP-cwt cells were infected with *S. aureus* 6850 mRFP. At 3 h p.i. phagosomal escape was determined by means of co-localization of mRFP and YFP signals. (C) For determination of intracellular replication of *S. aureus* 6850 GFP in HeLa and HAP1 cells the mean GFP fluorescence (AFU), which represents the bacterial load, was measured by flow cytometry. (D) Microscopic images of uninfected and *S. aureus* 6850 GFP infected HeLa and HAP1 cells 2 and 4 h p.i. (gray: phase contrast, green: *S. aureus*, scale bar: 50 μm). (E) HeLa and HAP1 cells infected with *S. aureus* 6850 were stained with the cell-impermeable dye 7-AAD at 5 h p.i. and analyzed by flow cytometry. (F) *S. aureus* 6850- or Cowan 1-infected HAP1 cells were lysed at 0, 1, 3, 4, 5 and 6 h p.i.. Uninfected cells or cells treated with 10 μg/ml cycloheximide (CHX) and 10 ng/ml TNFα for 7 h were used as control. Proteins were separated by SDS-PAGE and transferred onto a PVDF membrane. α-spectrin was detected using an anti-α-spectrin antibody and an anti-β-tubulin antibody served as loading control. (G, H) HAP1 R-Geco cells were infected with *S. aureus* 6850 GFP. (G) Stills of time-lapse imaging are shown (green: *S. aureus*, red: R-Geco, gray: brightfield, scale: 50 µm). (H) Relative R-Geco fluorescence was quantified over the time period of infection in single uninfected cells (ni) or cells infected with *S. aureus* 6850. (I) Immunoblot analysis of HAP1 CAPN1 KO and CAPN4 KO clones generated using CRISPR/Cas9 was performed to check gene knock-out. Assay was performed as described for (F) and antibodies against Calpain 1 and 4 and β-actin as loading control were used. Statistical significance was assayed using unpaired t-test.

**Video S1. Intracellular *S. aureus* induces a rise in cytosolic Ca**^**2+**^. HeLa R-Geco cells were infected with *S. aureus* 6850 and imaged over time to visualize intracellular Ca^2+^-concentrations (green: *S. aureus*, red: R-Geco, gray: brightfield).

**Video S2. *S. aureus* secreted factors induce cytosolic Ca**^**2+**^**-signaling and - overload**. HeLa R-Geco cells were treated with 10 % sterilized supernatant of a *S. aureus* 6850 overnight culture and imaged over time to visualize intracellular Ca^2+^- concentrations (red: R-Geco, gray: brightfield).

